# Flagellar perturbations activate adhesion through two distinct pathways in *Caulobacter crescentus*

**DOI:** 10.1101/2020.07.21.215269

**Authors:** David M. Hershey, Aretha Fiebig, Sean Crosson

**Affiliations:** Department of Microbiology and Molecular Genetics, Michigan State University, East Lansing, MI 48824, USA; Department of Biochemistry and Molecular Biology, University of Chicago, Chicago, IL 60637, USA

**Keywords:** Adhesion, flagellum, motility, holdfast, biofilm

## Abstract

Bacteria carry out sophisticated developmental programs to colonize exogenous surfaces. The rotary flagellum, a dynamic machine that drives motility, is a key regulator of surface colonization. The specific signals recognized by flagella and the pathways by which those signals are transduced to coordinate adhesion remain subjects of debate. Mutations that disrupt flagellar assembly in the dimorphic bacterium *Caulobacter crescentus* stimulate the production of a polysaccharide adhesin called the holdfast. Using a genome-wide phenotyping approach, we compared surface adhesion profiles in wild-type and flagellar mutant backgrounds of *C. crescentus*. We identified a diverse set of flagellar mutations that enhance adhesion by inducing a hyper-holdfast phenotype and discovered a second set of mutations that suppress this phenotype. Epistasis analysis of the *flagellar signaling suppressor* (*fss*) mutations demonstrated that the flagellum stimulates holdfast production via two genetically distinct pathways. The developmental regulator PleD contributes to holdfast induction in mutants disrupted at both early and late stages of flagellar assembly. Mutants disrupted at late stages of flagellar assembly, which assemble an intact rotor complex, induce holdfast production through an additional process that requires the MotAB stator and its associated diguanylate cyclase, DgcB. We have assigned a subset of the *fss* genes to either the stator- or *pleD*-dependent networks and characterized two previously unidentified motility genes that regulate holdfast production via the stator complex. We propose a model through which the flagellum integrates mechanical stimuli into the *C. crescentus* developmental program to coordinate adhesion.

**Importance:** Understanding how bacteria colonize solid surfaces is of significant clinical, industrial and ecological importance. In this study, we identified genes that are required for *Caulobacter crescentus* to activate surface attachment in response to signals from a macromolecular machine called the flagellum. Genes involved in transmitting information from the flagellum can be grouped into separate pathways, those that control the *C. crescentus* morphogenic program and those that are required for flagellar motility. Our results support a model in which a developmental and a mechanical signaling pathway operate in parallel downstream of the flagellum and converge to regulate adhesion. We conclude that the flagellum serves as a signaling hub by integrating internal and external cues to coordinate surface colonization and emphasize the role of signal integration in linking complex sets of environmental stimuli to individual behaviors.

## Introduction

For microorganisms, solid surfaces serve as sites of nutrient accumulation, gateways into host tissues and shelters from environmental stresses (1–3). To access surface-associated niches, bacteria deploy specialized programs for seeking, recognizing and colonizing objects in their surroundings (4). These programs culminate in a pronounced transition away from a free-living, exploratory state and toward an adherent, sessile lifestyle (5–7). Sophisticated signaling networks that integrate a host of environmental cues are used to coordinate the motile-to-sessile switch (8–10). The complexity of these circuits reflects the perilous nature of committing to colonization programs under sub-optimal conditions.

A transenvelope machine called the flagellum drives cellular motility and plays a critical role at numerous stages of surface colonization(11, 12). The flagellum is synthesized in a stepwise process that is controlled by a transcriptional hierarchy(13). Assembly begins with the expression of class II genes that code for a rotor and secretion subcomplex that are inserted in the cytoplasmic membrane(14, 15). Upon completion of the class II program, assembly proceeds outward with the incorporation of an envelope spanning basal body (class III genes) followed by the secretion of an extracellular filament (class IV genes) (16, 17). Stator subcomplexes that surround the rotor utilize ion gradients across the cytoplasmic membrane to generate torque by turning the hook-basal body complex and its associated filament, propelling the cell forward (Fig 1) (18, 19). Flagellar motors are highly attuned to environmental conditions. They support motility under diverse conditions, modulate torque in response to changing loads and alter rotational bias to support complex swimming patterns(20–23).

**Figure 1.**
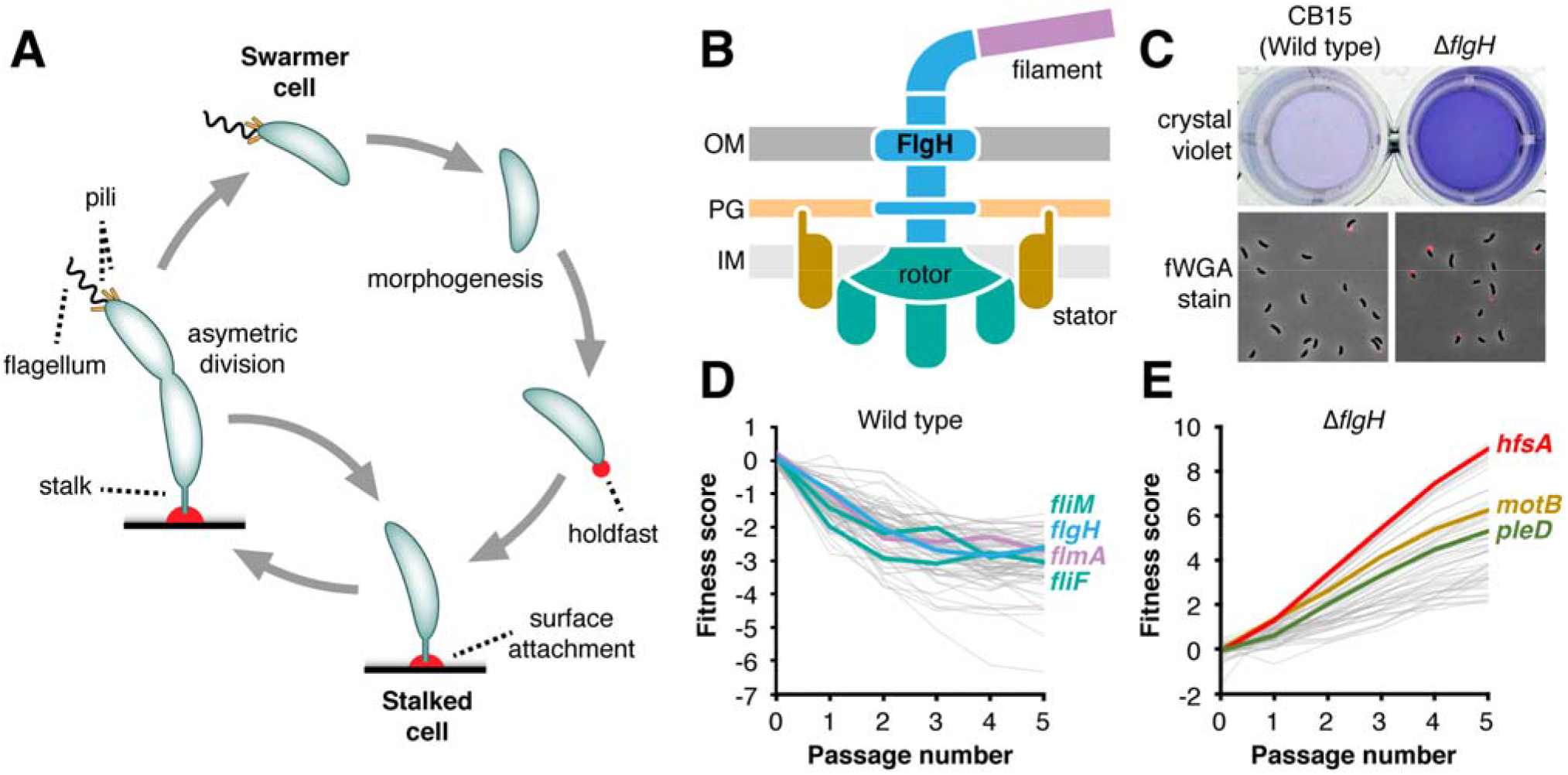
Identifying genes that link the flagellum to holdfast production. A) The asymmetric division cycle of *C. crescentus*. Sessile stalked cells divide to release a newborn swarmer cell that displays a flagellum and type IV pili. Quiescent swarmer cells undergo a morphological transition to become replication competent stalked cells. Transitioning swarmer cells can make an adhesin called the holdfast (red) that promotes surface attachment. B) Schematic of flagellar architecture. A central hook-basal body (HBB) complex (blue) spanning the cell envelope tethers a long extracellular filament (purple) to the surface of the cell. Multiple stator subunits (gold) that surround the inner membrane embedded rotor (teal) use ion translocation to turn the HBB and its associated filament. Outer membrane (OM), peptidoglycan (PG) and inner membrane (IM) layers of the envelope are shown. C) The Δ*flgH* mutant shows increased surface colonization (top), as measured by crystal violet (CV) staining, and a higher proportion of holdfast producing cells relative to wild type (bottom). Holdfasts were stained with Alexa 594-wheat germ agglutinin (fWGA). D) Hyper-adhesive mutants identified by adhesion profiling in defined medium. The 75 genes with the strongest hyper-adhesive profiles are plotted, and specific flagellar assembly genes are highlighted. Colors correspond to the structural proteins depicted in panel B. The plotted genes are listed in Table S1. E) The *flagellar signaling suppressor* (*fss*) genes identified by adhesion profiling in the Δ*flgH* background. The 50 genes with the strongest contributions to adhesion in the Δ*flgH* background are plotted. *hfsA*, a gene required for holdfast biosynthesis, is highlighted along with two *fss* genes, *pleD* and *motB*. The plotted genes are listed in Table S2. For panels D and E, each line represents the average fitness values for a single gene plotted as a function of time in the sequential passaging experiment. Hyper-adhesive mutants are depleted more rapidly than neutral mutants during selection in cheesecloth. Mutated genes (*fli,flg, flm*, etc.) that display increased attachment to cheesecloth show steadily decreasing fitness scores as a function of passage number. Mutants with reduced adhesion are enriched in broth when grown with cheesecloth. Mutated genes that display decreased adhesion (*hfs* and *fss*) show steadily increasing fitness scores.

Paradoxically, flagellar motility must be repressed during sessile growth but is also required for efficient surface colonization(6, 24–26). During the initial stages of attachment, swimming is thought to promote productive interactions with target substrates by providing energy needed to overcome repulsive forces at the liquid-solid interface(27). The flagellum also plays an additional regulatory role in activating the motile-to-sessile transition by recognizing physical contact with solid substrates(11). Such tactile sensing events serve as critical cues for initiating colonization programs, but the mechanistic basis for how bacteria sense and respond to physical stimuli remains controversial.

The dimorphic bacterium *Caulobacter crescentus* is uniquely adapted to surface colonization. Cell division in *C. crescentus* is asymmetric and yields to two distinct cell types(28). Newborn swarmer cells are flagellated, produce type IV pili (T4P) and cannot initiate replication (29, 30). These motile cells undergo a morphogenic transition to become replication-competent stalked cells by replacing their flagellum and pili with a specialized envelope extension called the stalk(31). During the swarmer-to-stalked transition, *C. crescentus* can produce a polysaccharide adhesin called the holdfast that is displayed at the tip of the stalked cell where it promotes attachment to surfaces (Fig 1) (7, 32). Holdfast production is the primary determinant of surface colonization in *C. crescentus*, and its regulation is elaborate (26, 33). In addition to cell cycle cues(7, 34), nutrient availability(35), light(36) and redox status(37), mechanical contact has been implicated as an important activator of holdfast assembly (38). Recent evidence suggests that both flagella and T4P can stimulate holdfast production in response to contact with a surface(39, 40), but conflicting models have emerged for how these transenvelope machines survey and disseminate mechanical information(34, 41).

Here, we used an unbiased phenotyping approach called adhesion profiling to show that a diverse set of flagellar mutations induce a hyper-holdfast phenotype and to identify dozens of *<u>f</u>lagellar <u>s</u>ignaling <u>s</u>uppressor* (*fss*) genes that contribute to holdfast stimulation downstream of the flagellum, *fss* mutations suppress the hyper-adhesive effects of flagellar disruption through two distinct pathways. Select regulators of cell-cycle progression are involved in stimulating adhesion upon flagellar disruption, while components of the stator subcomplexes contribute to holdfast stimulation specifically in mutants that can assemble an inner membrane rotor. We assigned roles for two previously uncharacterized genes roles in the stator-dependent pathway and demonstrated that they promote the ability of the stator subunits to turn the flagellar filament. Our results provide new insight into load sensing by the *C. crescentus* motor and highlight a novel link between flagellar assembly and morphogenesis. We propose a broad role for the flagellum in coordinating cellular physiology through its role as a signaling hub that integrates internal and external cues.

## Results

### A complex gene network links the flagellum to holdfast production

We previously described a method called adhesion profiling by which a barcoded transposon library is sequentially passaged in the presence of a cellulose-based substrate. Adhesive mutants become depleted as they colonize the substrate, enriching for mutants with attachment defects in the surrounding broth. By monitoring the mutant population over time, we quantified each gene’s contribution to adhesion at the genome scale (33). This initial study identified a set of hyper-adhesive mutants that included genes involved in flagellar assembly, which suggested the presence of a specific signaling pathway linking cues from the flagellum to holdfast production (Fig 1C). We modified our genetic selection to identify a broader range of adhesion-activating mutations by using a defined medium (M2X) in which holdfast production is almost entirely repressed in wild-type *C. crescentus*(35). Under these conditions, dozens of genes displayed adhesion profiles indicative of hyper-adhesion (Fig 1D and Table S1). Though numerous functional categories were represented in this gene set, the overwhelming majority of hyper-adhesive phenotypes were observed in mutants with predicted disruptions to flagellar assembly, chemotaxis or other flagellar processes.

We focused on the holdfast phenotype for a mutant (Δ*flgH*) lacking the gene for the flagellar L ring protein(42) growing in M2X medium (Fig 1C). Consistent with previous reports (33, 34), crystal violet (CV) staining of surface attached cells was elevated in Δ*flgH* cultures relative to wild type, and a larger proportion of cells displayed a holdfast when stained with fluorescently labelled wheat germ agglutinin (fWGA, Fig 1C). Mutating genes that code for extracellular components of the *C. crescentus* flagellum was proposed to increase adhesion by rendering cells hyper-sensitive to surface contact(40), but our results indicated that the Δ*flgH* mutant displayed elevated holdfast production when grown in liquid without an activating surface. Though our standard fWGA staining protocol includes brief centrifugation steps, we confirmed that the proportion of holdfast producing cells did not change when centrifugation was omitted and cells were imaged directly from liquid cultures (Fig S1). Additionally, we found that the Δ*flgH* mutant released holdfast polysaccharide directly into spent liquid medium (Fig S1), another hallmark of surface-independent holdfast activation (43). These results are inconsistent with the model that the Δ*flgH* mutant is hyper-sensitive to surface contact. Instead, elevated adhesion in Δ*flgH* results from surface contact-independent increases in both the proportion of cells that assemble a holdfast and the amount of secreted holdfast polysaccharide. We conclude that flagellar mutations act as gain of function activators of holdfast production.

To dissect potential pathways linking the flagellum to holdfast production, we constructed a barcoded Tn-*Himar1* library in a Δ*flgH* background and performed a second adhesion profiling experiment with the goal of identifying mutations that suppress the hyper-holdfast phenotype. As in wild type, genes required for holdfast synthesis (*hfs*) were the strongest determinants of adhesion in the Δ*flgH* mutant. In addition, we identified a few dozen genes (called *fss* for flagellar signaling suppressor) that contribute to adhesion specifically in the Δ*flgH* background (Fig 1E, Table S2). While, many of the *fss* genes are uncharacterized, insertions in genes known to promote flagellar rotation, chemotaxis, cell cycle progression and other physiological processes had *fss* phenotypes as well. Both the abundance and functional diversity of the suppressors point to a complex signaling network that links adhesion to flagellar motility.

### Distinct adhesion patterns in flagellar assembly mutants

Two of the *fss* genes, *motB* and *pleD*, are known to regulate holdfast production under specific conditions. *motB*, which codes for one of the flagellar stator proteins, is required for rapid holdfast synthesis after surface contact in microfluidic chambers(40). *pleD*, which codes for a diguanylate cyclase that regulates morphogenesis during the swarmer-to-stalked transition(44), contributes to increased holdfast production in a flagellar hook mutant background through a process independent of surface contact(34). Previous examinations of these two genes have produced conflicting models for how surface contact, flagellar assembly and filament rotation modulate holdfast production. Identifying mutations in both *pleD* and *motB* as suppressors of Δ*flgH* suggested that we could clarify the signaling pathway that links flagellar perturbations to holdfast production.

We used CV staining to examine how disrupting *pleD* and *motB* affects adhesion to polystyrene in various flagellar mutant backgrounds. Disrupting the early stages of flagellar assembly by deleting the class II genes *fliF* or *fliM* led to a hyper-adhesive phenotype that was strongly suppressed by deletion of *pleD* but that was not affected by deletion of *motB*. In contrast, when holdfast production was stimulated by deletion of the class III gene *flgH* or disruption of flagellin secretion (Δ*flmA*)(45), the hyper-adhesive phenotype was suppressed by introducing either a *pleD* or a *motB* deletion (Fig 2A). Thus, flagellar mutants stimulate adhesion through different mechanisms. Mutants that disrupt the early stages of assembly activate holdfast production through *pleD*, while mutants in which assembly is stalled at later stages stimulate adhesion through both *pleD* and *motB*.

**Figure 2.**
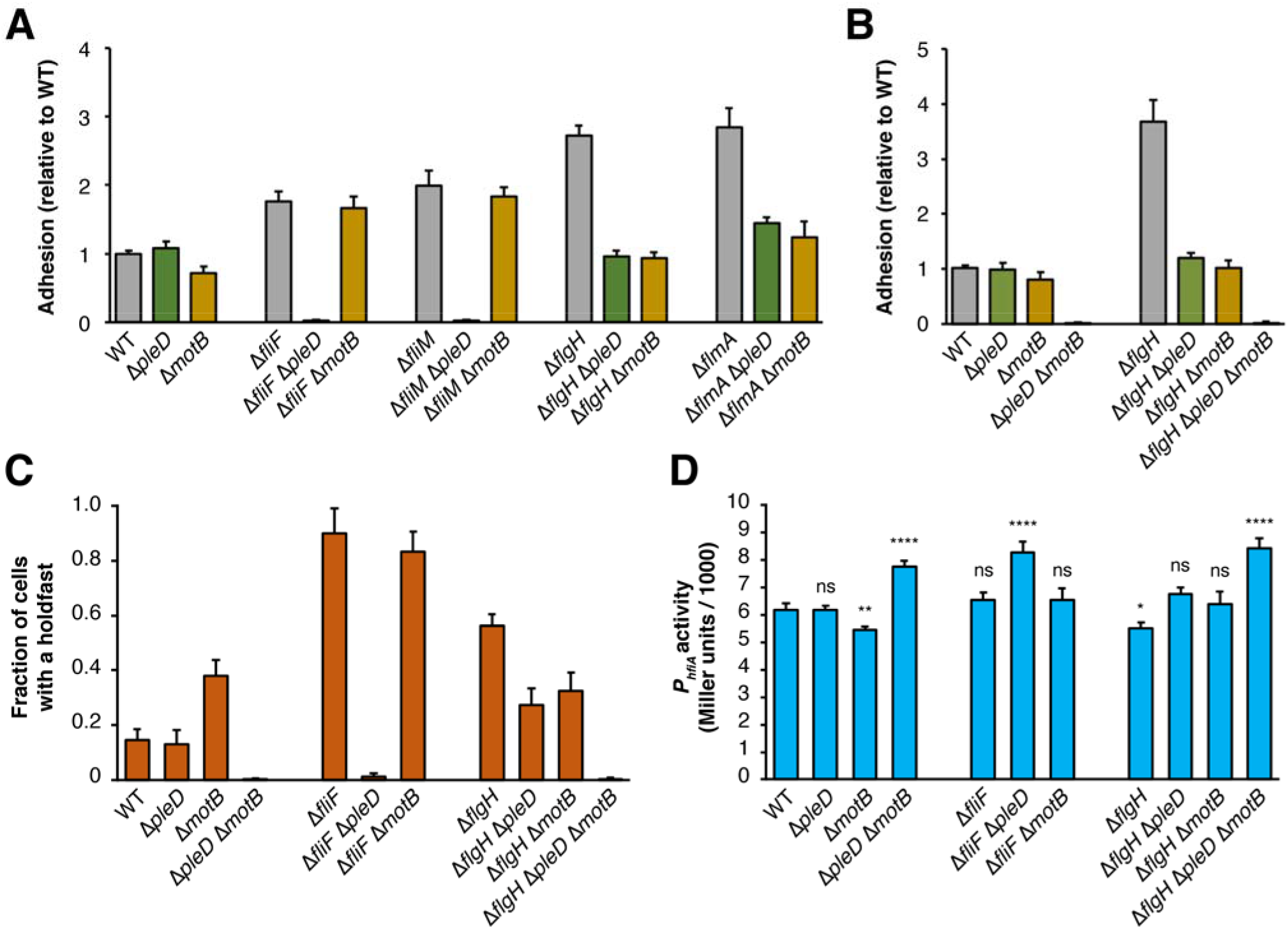
Two distinct signaling pathways operate downstream of the flagellum to control adhesion. A) Crystal violet-based attachment assay showing suppression of the hyper-adhesive phenotypes by *pleD* and *motB* in early (Δ*fliF* and Δ*fliM*) and late (Δ*flgH* and Δ*flmA*) flagellar assembly mutants. Mean values from six biological replicates are shown with error bars representing the associated standard deviations. B) Crystal violet-based attachment assay showing the additive effects of Δ*motB* and Δ*pleD* on adhesion in the WT and Δ*flgH* backgrounds. Mean values from five biological replicates are shown with the associated standard deviations. C) Fraction of cells with a holdfast in flagellar mutant and suppressor backgrounds. Holdfasts were stained and counted from log phase cultures as described in Materials and Methods. D) *hfiA* transcription in flagellar mutant and suppressor backgrounds as measured by beta-galactosidase activity from a *P_hfiA_-lacZ* reporter. Mean values from three biological replicates collected on two separate dates for a total of six replicates are shown with the associated standard deviations. Statistical significance was evaluated with ANOVA followed by Tukey’s multiple comparison test. Significance compared to wild-type is indicated above each bar. (ns): not significant; (*): P < 0.1; (**): P < 0.01; (****); P < 0.0001. A full statistical analysis of the CV staining, holdfast count and LacZ activity measurements is reported in Table S3.

### Two pathways modulate holdfast production downstream of the flagellum

The distinct suppression patterns in *pleD* and *motB* mutants indicated that multiple pathways function downstream of the flagellum to influence adhesion. Indeed, combining the Δ*pleD* and Δ*motB* mutations reduced CV staining to near un-detectable levels in both the wild-type and Δ*flgH* backgrounds, supporting the model that *pleD* and *motB* control attachment through distinct mechanisms (Fig 2B, Fig S2A). The severe adhesion defect observed for the Δ*pleD* Δ*motB* double mutant demonstrates that the *pleD*- and *motB*-dependent pathways do not operate exclusively in the context of flagellar mutants. Either the *pleD* or *motB* pathway must be intact for *C. crescentus* to colonize surfaces.

We quantified holdfast production by staining cells from a representative panel of mutants with fWGA. The proportion of cells displaying a holdfast was elevated in both early (Δ*fliF*) and late (Δ*flgH*) flagellar assembly mutants, and the suppression patterns seen by CV staining were recapitulated with fWGA staining. Holdfast production was elevated to similar levels in Δ*fliF* and Δ*fliF* Δ*motB* but nearly eliminated in Δ*fliF* Δ*pleD*. Introducing either the Δ*pleD* or the Δ*motB* mutations reduced holdfast production in a Δ*flgH* background, and holdfast production was nearly undetectable in Δ*pleD* Δ*motB* and Δ*flgH* Δ*pleD* Δ*motB* cultures (Fig 2C and Table S3). We did identify modest discrepancies between the holdfast production and polystyrene colonization measurements. Although surface attachment was indistinguishable from wild type in Δ*motB* cultures, holdfast production in this mutant was elevated. This agrees with previous measurements indicating that non-motile strains display holdfast-independent surface colonization defects (33, 34). Separately, CV staining was higher in Δ*flgH* cultures than in Δ*fliF* cultures, but the proportion of holdfast producing cells was higher in the Δ*fliF* mutant. Because both strains are non-motile, the discrepancy is likely due to modulation of additional holdfast-independent colonization factors such as type IV pilus dynamics(33, 46).

Finally, we examined expression of the *holdfast inhibitor A* (*hfiA*) gene, a key regulator that inhibits adhesion by targeting a glycosyltransferase in the holdfast biosynthesis pathway (Fig 2D)(35). Increased holdfast production in the Δ*flgH* and Δ*motB* backgrounds is accompanied by a decrease in *P_hifA_-lacZ* reporter activity, but elevated holdfast production in the Δ*fliF* mutant occurs without a reduction in *hfiA* transcription (Fig 2D and Table S3). Introducing either the Δ*motB* or Δ*pleD* mutations into the Δ*flgH* background restored *P_hifA_* activity to wild-type levels. These measurements show that *pleD* is required for downregulation of *hfiA* in the Δ*flgH* background, but do not clarify the role of *motB*. It remains unclear how the Δ*motB* mutation attenuates *hfiA* promoter activity in the wild-type background but restores normal expression in a Δ*flgH* mutant. Transcription from the *hfiA* promoter was elevated in the Δ*pleD* Δ*motB*, the Δ*flgH* Δ*pleD* Δ*motB* and the Δ*fliF* Δ*pleD* mutants, indicating that activation of *hfiA* expression contributes to the severe holdfast production defect in these three strains.

Transcription of *hfiA* is finely tuned by a complex hierarchy of transcription factors such that small changes in expression have significant impacts on holdfast production (33, 35, 37). The three non-adhesive mutants analyzed in Figure 2 display robust increases in *hfiA* expression, and the Δ*fliF* mutant shows a striking increase in holdfast production that clearly occurs independently of *hfiA* regulation. However, *P_hifA_-lacZ* activity differences for other key strains are modest and do not correlate perfectly with direct measurements of holdfast production. While cell-cycle control and post-transcriptional processes can be masked in bulk reporter measurements, the expression level changes in flagellar signaling mutants are less pronounced than for regulatory systems that target *hfiA* directly (35, 37). A significant portion of adhesion control exerted by the flagellum likely occurs independent of *hfiA* regulation.

### Parsing regulatory networks with epistasis analysis

The distinct activation profiles observed in early and late flagellar assembly mutants were used to assign *fss* genes to either the *pleD*- or *motB*-dependent signaling pathways. We predicted that *pleD* and other genes involved in stalked cell morphogenesis make up a “developmental” signaling pathway and that genes associated with stator activity make up a second, “mechanical” pathway. This model predicts that developmental pathway mutants should block holdfast stimulation in both early and late flagellar assembly mutant backgrounds, while mechanical pathway mutants should suppress hyper-adhesion specifically in late assembly mutants. Two additional *fss* genes, *shkA* and *dgcB*, were used to test these predictions. *shkA* encodes a histidine kinase that regulates stalk development(47), and *dgcB* codes for a diguanylate cyclase that physically associates with stator subcomplexes(40). Deleting *shkA* suppressed the hyper-adhesive effects of both the Δ*fliF* (early) and the Δ*flgH* (late) mutations, while deleting *dgcB* suppressed adhesion in the Δ*flgH* background but had no effect in the Δ*fliF* background (Fig 3A). Furthermore, adhesion was nearly eliminated when the Δ*shkA* mutation was introduced into the Δ*motB* or Δ*dgcB* backgrounds (Fig S2), confirming that *shkA* signals through a mechanism distinct from that of mechanical pathway genes. These results provide further support for a model in which both a developmental pathway associated with stalked cell morphogenesis and a mechanical pathway associated with stator activity function downstream of the flagellum to activate adhesion.

**Figure 3.**
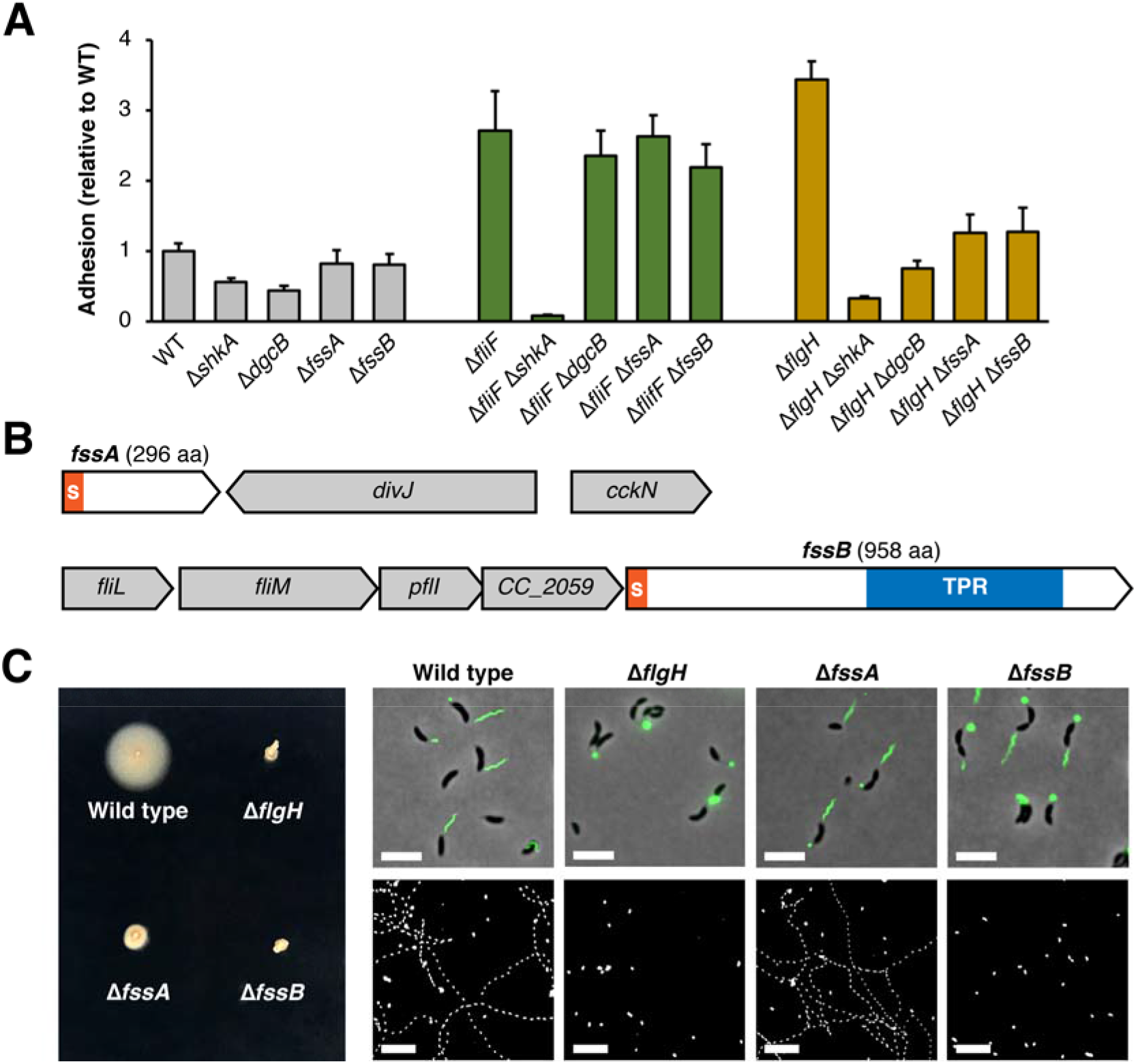
Two novel motility genes contribute to activation of the mechanical pathway. A) Crystal violet-based attachment assay evaluating suppression of early and late flagellar assembly mutants by individual *fss* genes. Mean values from six biological replicates are shown with error bars representing the associated standard deviations. B) Genomic context of the *fssA* and *fssB* genes. Orange S: secretion signal; blue bar: tetratricopeptide repeat region. C) Motility phenotypes of the Δ*fssA* and Δ*fssB* mutants. Left: soft-agar motility assay. Top right: flagellar filaments stained with Alexa488-maleimide after introduction of the *fljK^T103C^* allele into the indicated mutants. Note that the maleimide dye cross-reacts with holdfast. Scale bars represent 5μm. Bottom right: maximum projections from time-lapse microscopy. Cells appear in white, and tracks for motile cells appear as dotted lines. Scale bars represent 25μm.

We also used epistasis to place two uncharacterized genes identified as Δ*flgH* suppressors into the mechanical signaling pathway. *fssA* (*CC_1064; CCNA_01117*) encodes a protein with a Sec/SP1 secretion signal and no predicted functional domains. *fssB* (*CC_2058; CCNA_02137*) encodes a protein with a Sec/SP1 secretion signal and a predicted tetratricopeptide repeat (TPR) domain. Deleting either *fssA* or *fssB* did not affect adhesion in the wild-type or Δ*fliF* backgrounds but suppressed the hyper-adhesive phenotype in Δ*flgH* (Fig 3A), providing evidence that *fssA* and *fssB* contribute to holdfast stimulation through the stator-dependent, mechanical pathway.

### New motility factors contribute to mechanical activation

To understand how *fssA* and *fssB* regulate holdfast synthesis, we examined the motility phenotypes of Δ*fssA* and Δ*fssB* deletions. Both mutants were severely impaired in their ability to spread through soft agar. When a flagellin allele (*fljK^T103C^*) coding for an FljK protein that can be stained with maleimide-conjugated dyes(34) was introduced, flagellar filaments were observed in both Δ*fssA* and Δ*fssB* cells. Thus, the motility phenotypes in these mutants are not caused by disruptions to flagellar assembly. Examination of individual cells in liquid broth revealed that Δ*fssB* cells were non-motile, while some Δ*fssA* cells retained the ability to swim (Fig 3C). Thus, the Δ*fssB* mutant displays a paralyzed flagellum phenotype analogous to a Δ*motB* mutant, but the motility phenotype in Δ*fssA* is specific to soft agar.

Over the course of our studies, we observed that the Δ*fssA* mutant had a propensity to begin spreading through soft-agar after prolonged incubation on plates (Fig 4A). Colonies migrated from the inoculation site in an anisotropic manner, suggesting that second-site suppressors of the motility defect had emerged. Indeed, single colonies isolated from motile Δ*fssA* flares were indistinguishable from wild-type when re-inoculated into soft-agar. Fourteen of these motile suppressors were analyzed by whole genome sequencing to identify the causative mutations. Each isolate harbored a missense mutation in one of the stator genes. Three contained a mutation in *motA*, and eleven contained a mutation in *motB*. Nine of the eleven *motB* mutations disrupt the same residue, serine 52, and six produce the same allele, *motB^S52C^* (Fig S3).

**Figure 4.**
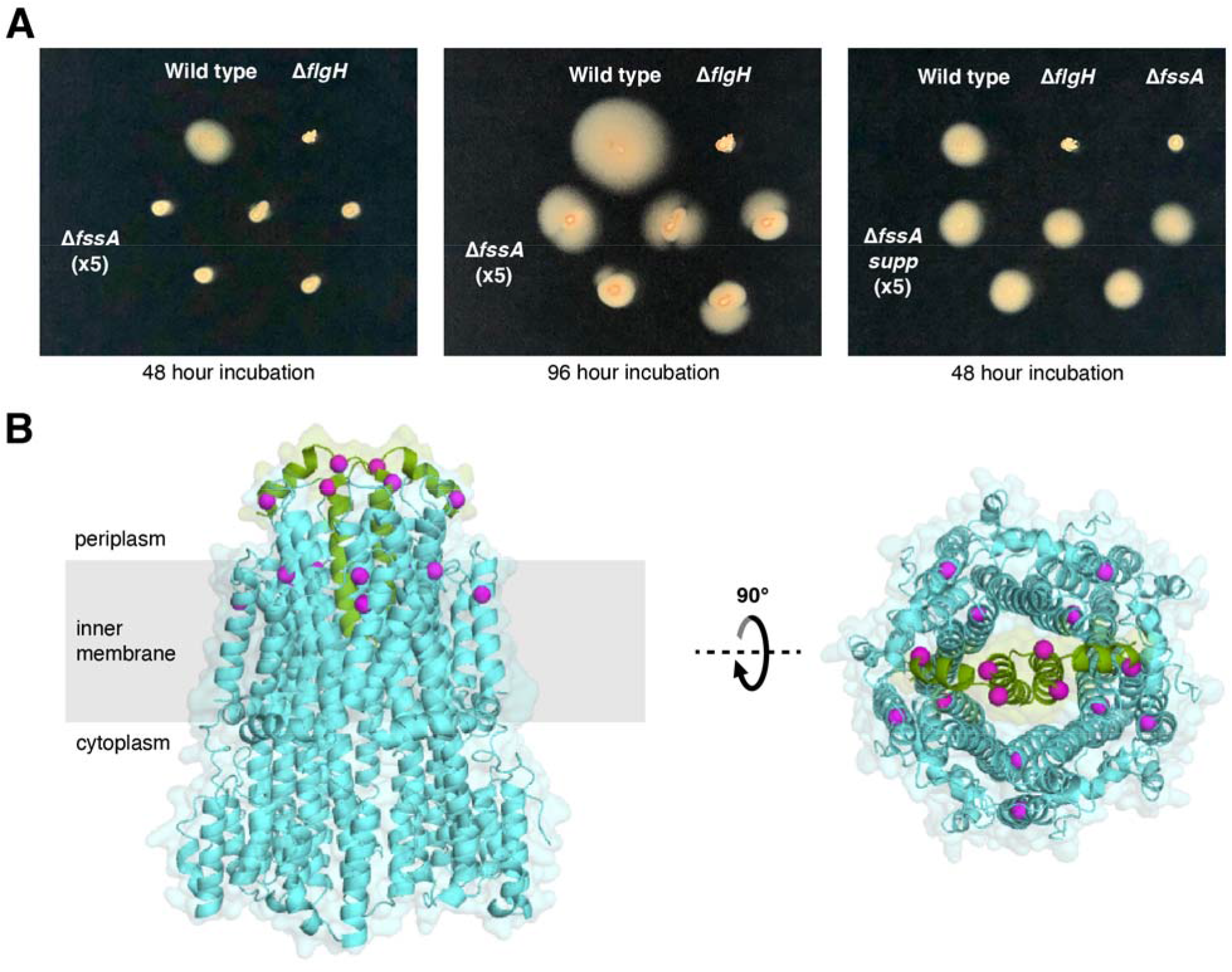
Suppression of the ΔfssA motility phenotype by second site mutations in the stator genes. A) Soft-agar motility assays showing the emergence of motile flares after prolonged incubation of the Δ*fssA* mutant. Single colonies isolated from the leading edge of flares (middle image) displayed wild-type motility when reinoculated into soft-agar (right image). B) Mapping of Δ*fssA* suppressors onto a homology model of the *C. crescentus* stator. Mutations are located at the periplasmic face of the complex. Identification of multiple mutations in the “plug” region of MotB suggests that the suppressing mutations activate ion translocation ectopically.

We used a cryoelectron microscopy reconstruction of the MotAB stator from *Campylobacter jejuni*(48) to predict the structure of the *C. crescentus* stator complex. The resulting homology model contains a characteristic transmembrane channel composed of five MotA subunits that is capped at its periplasmic face by two MotB protomers(48, 49). When Δ*fssA* suppressing mutations were mapped onto this model they displayed a clear bias toward residues on the periplasmic face of the complex (Fig 4B), with the *motB* mutations all disrupting a region known as the plug. Deleting the plug allows ion translocation through the stators in the absence of productive engagement with a rotor(50), and missense mutations in the plug have been shown to support motility under nonpermissive conditions through gain of function activation of the motor(51).

The ability of plug mutations to suppress the Δ*fssA* motility defect indicates that Δ*fssA* and Δ*fssB* display related motility phenotypes. Δ*fssB* produces an inactive flagellar motor that cannot turn a filament (Fig 3C), while Δ*fssA* assembles a modestly defective motor that supports full motility in soft-agar only when the stators are activated by mutations predicted to increase ion translocation (Fig 4). A Δ*fssA* Δ*fssB* double mutant did not spread through soft-agar even after prolonged incubation, confirming that the lack of motor rotation in Δ*fssB* is epistatic to the partial defect in Δ*fssA* (Fig S3). Furthermore, our data indicate that *fssA* and *fssB* support flagellar signaling by the same the mechanism, as the Δ*fssA* and Δ*fssB* mutations did suppress Δ*flgH* hyper-adhesion in an additive manner (Fig S3). We conclude that *fssA* and *fssB* are required for proper stator activity in *C. crescentus. We* propose that mutating either gene disrupts both the stator’s ability to promote motility and its capacity to transduce mechanical signals.

### Separate mechanisms for activating c-di-GMP production

Though the developmental and mechanical pathways can be separated genetically, they ultimately converge to modulate holdfast production. Each pathway includes a diguanylate cyclase predicted to synthesize bis-(3’-5’)-cyclic diguanosine monophosphate (c-di-GMP), a second messenger that promotes surface-associated behaviors in bacteria(52). In *C. crescentus*, c-di-GMP binds numerous downstream effectors to activate stalk assembly(53), cell cycle progression(54, 55) and holdfast synthesis (40, 56). To test the role of c-di-GMP synthesis in linking cues from the flagellum to holdfast production, we examined catalytically inactive alleles of *pleD* and *dgcB*. In contrast to wild-type alleles, *pleD^E370Q^*(57) and *dgcB^E261Q^*(40) failed to restore hyper-adhesion in the Δ*flgH* Δ*pleD* and Δ*flgH* Δ*dgcB* backgrounds, respectively, confirming that c-di-GMP synthesis by these enzymes is required to support flagellar signaling through both the mechanical and developmental pathways (Fig 5A).

**Figure 5.**
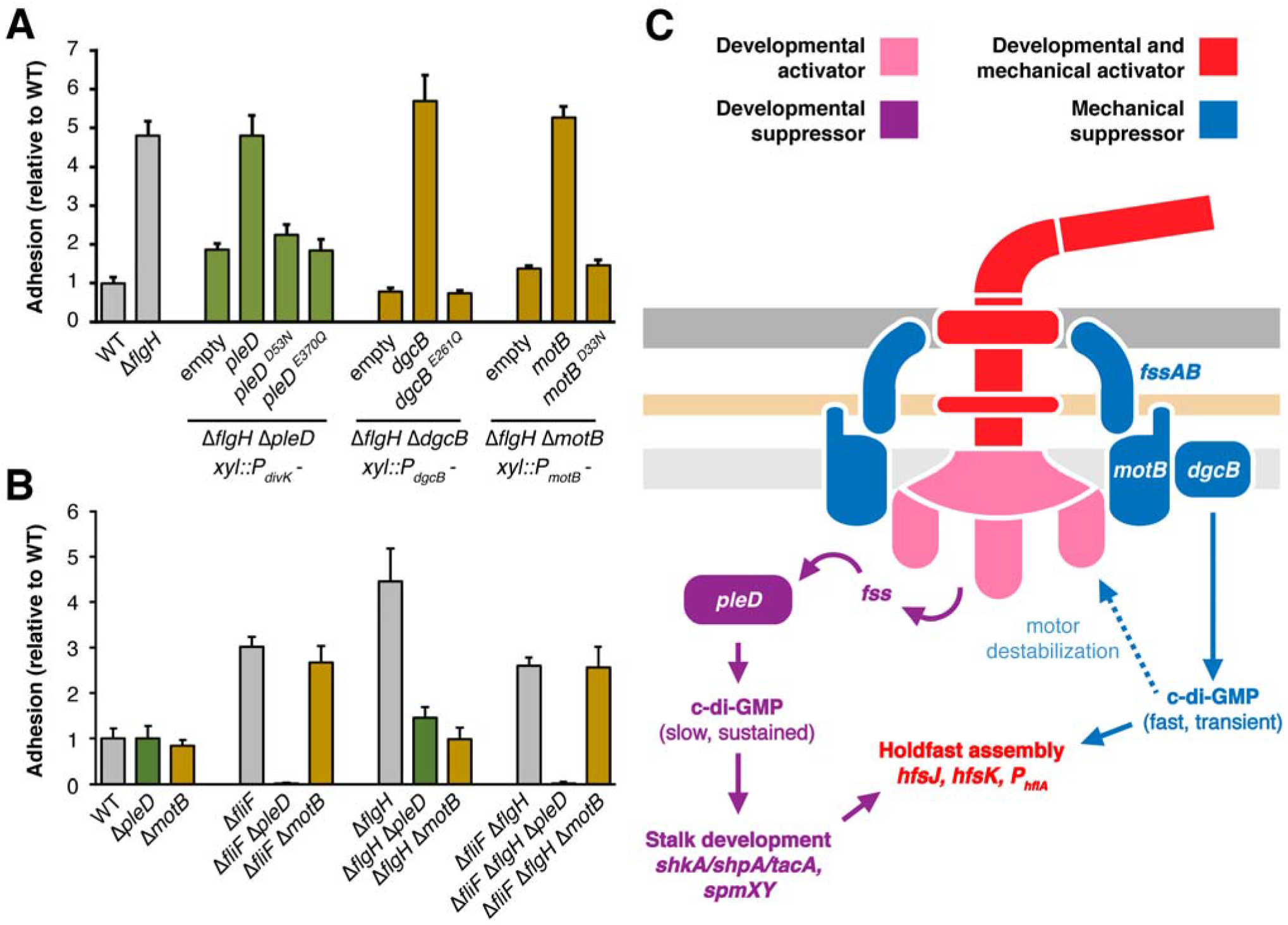
Convergence of flagellar signaling on cyclic-di-GMP. A) Crystal violet-based attachment assay showing that inactive alleles of *fss* genes in the developmental (*pleD*) and mechanical (*dgcB* and *motB*) pathways do not support flagellar signaling. Each allele is expressed from the gene’s native promoter and integrated as a single copy at the *xylX* locus. The genotype and promoter for each strain are indicated below the solid line. Mean values from seven biological replicates are shown with error bars representing the associated standard deviations. B) Crystal violet-based attachment assay showing that early flagellar mutants are epistatic to late flagellar mutants for activation of the mechanical pathway. The Δ*fliF* Δ*flgH* strain phenocopies the Δ*fliF* strain and demonstrates that the effect of *motB* on adhesion requires early stages of flagellar assembly. Mean values from five biological replicates are shown with the associated standard deviations. C) Two signaling pathways operate downstream of the flagellum in *C. crescentus*. We propose that mechanical signals are transmitted through the motor to activate DgcB, producing a transient burst of c-di-GMP synthesis that directly activates holdfast biosynthesis enzymes. Persistent filament obstruction activates PleD by destabilizing the motor and triggering flagellar disassembly. PleD activation through this mechanism is predicted to induce sustained c-di-GMP production and initiation of the transcriptional program that leads to stalk development.

We examined mechanisms by which the diguanylate cyclase activities of PleD and DgcB are activated during flagellar signaling. PleD contains a receiver domain at its N-terminus that includes a canonical aspartyl phosphorylation site(57, 58). Introducing the non-phosphrylatable *pleD^D53N^* allele failed to restore hyper-adhesion in the Δ*flgH* Δ*pleD* background (Fig 5A). We conclude that phosphorylation of the PleD receiver domain is required for flagellar perturbations to stimulate holdfast production through the developmental pathway. Proton translocation by MotAB stators is used to generate torque for flagellar filament rotation. A *motB* allele (*motB^D33N^*) that prevents proton flux through the stator(40, 59) does not support flagellar signaling in the Δ*flgH* background (Fig 5A). We conclude that proton translocation through MotB is required for flagellar perturbations to stimulate holdfast production via the mechanical pathway.

Our data indicate that active proton translocation is required for mechanical stimulation. This model provides a possible explanation for the disparate suppression patterns we observed in early and late flagellar assembly mutants. We predicted that the mechanical pathway is inactive in early flagellar mutants because stator subunits cannot engage with the motor when rotor assembly is incomplete. Indeed, Δ*fliF* was epistatic to Δ*flgH* with respect to suppression by Δ*pleD* and Δ*motB* (Fig 5B). Adhesion in the Δ*fliF* Δ*flgH* mutant was eliminated when *pleD* was deleted but remained unchanged when the Δ*motB* mutation was introduced, supporting the model that inner membrane rotor assembly is required for activation of the mechanical pathway.

## Discussion

Many bacteria alter their behavior after contact with exogenous surfaces, and flagellar motility is a key regulatory determinant of these responses(38, 60–63). However, efforts to dissect contact-dependent signaling pathways have been confounded by contributions from multiple mechanosensors(62, 64), a reliance on non-canonical signaling machinery(9, 40) and the prevalence of transcription-independent responses(7, 65). In this study, we leveraged the hyper-adhesive phenotype induced upon mutation of flagellar assembly genes to dissect the genetic basis for adhesion control by the flagellum in *C. crescentus*. We used a high-throughput phenotyping approach to identify mutations that stimulate adhesion and to classify a large group of genes called *flagellar signaling suppressors* (*fss*) that contribute to increased holdfast production when flagellar assembly is disrupted. The results have clarified important features of how the *C. crescentus* flagellum regulates adhesion and provided a framework for disentangling signaling networks that control bacterial behavior.

Two genes identified in the *fss* screen, *pleD* and *motB*, have been previously shown to link flagellar function to adhesion, but conflicting models were proposed for how these genes regulate holdfast production(34, 40). We showed that *pleD* and *motB* participate in genetically distinct pathways for activating adhesion. *pleD* and its downstream effector *shkA* contribute to increased holdfast production when any stage of flagellar assembly is disrupted. *motB*, the gene for its associated diguanylate cyclase, *dgcB*, and two previously uncharacterized motility genes contribute to adhesion specifically in late flagellar mutants that retain the ability to assemble inner membrane rotors. We conclude that a mechanical pathway and a developmental pathway operate in parallel to link flagellar function to holdfast production.

Strains harboring deletions (Δ*motB*, Δ*fssA*, or Δ*fssB*) or mutant alleles (*motB^D33N^*) that disrupt motility without affecting filament assembly cannot support activation of the mechanical pathway (Figs 2A, 3A and 5A). Additionally, early flagellar assembly mutants display epistatic effects on late assembly mutants by eliminating *motB*’s involvement in stimulating adhesion (Fig 5B). These results suggest that blocking inner membrane rotor (MS- and C-ring) assembly subverts the mechanical pathway by preventing stator engagement and help to explain the range of behavioral effects often observed in flagellar assembly mutants(5, 40, 66). We conclude that intact motors capable of generating torque are required for mechanical activation of the *C. cresecentus* flagellum. Increased load on rotating filaments leads to the recruitment of additional stators to the motor in enteric bacteria(67, 68), and a similar resistance sensing mechanism likely supports tactile sensing in *C. crescentus*. Mutants in mechanical pathway genes such as *fssA* and *fssB* should prove useful in describing the structural basis for how such changes in load are sensed by the flagellar motor.

Key features of the mechanical pathway mirror the tactile sensing event described by Hug and colleagues(40), but certain conclusions must be reevaluated in light of our results. We showed that late flagellar mutants display increased holdfast production in M2X liquid medium (Fig 2C and Fig S1) in the absence of a surface and under nutrient conditions for which tactile sensing does not normally occur(34). We conclude that mutants lacking the outer parts of the flagellum do not show a hyper-sensitive surface response. Instead, we infer that the motor responds to the absence of a filament as it would to an obstructed filament by activating stator-dependent signaling ectopically. Consistent with this interpretation, preventing stators from productively engaging with the rotor, either by disrupting proteins required for stator function (Figs 2A and 3A) or by mutating early flagellar genes that code for rotor components, blocks mechanical signaling in late assembly mutants (Fig 5B). This explains the perplexing epistasis of inner parts of the flagellum over outer parts(69) and supports established models for tactile sensing through increased load on the filament(60) rather than the motor acting as a tetherless sensor(40). More broadly, the disparate phenotypes identified here for stator, rotor and hook-basal body mutants reflect an emerging pattern seen in other organisms and argue that signal bifurcation by the flagellum is a conserved feature throughout bacteria(70, 71).

The identification of a second, developmental pathway downstream of the flagellum is consistent with previous studies showing that late flagellar assembly mutants display contact-independent increases in holdfast production(33, 34). In fact, disrupting flagellar assembly at any stage stimulates adhesion (Fig 2A), and our analysis of this process highlights an overlooked role for the flagellum in controlling the *C. crescentus* developmental program. We explicitly characterized *pleD* and *shkA*, but other developmental regulators identified in the *fss* screen (*shpA, tacA*(47), *spmX*(72), *spmY*(73), *zitP, cpaM*(74), *sciP*(75), *rpoN*(76)) likely also act downstream of both early and late flagellar mutants to stimulate adhesion. Most of these genes influence flagellar assembly either directly or by altering cell cycle progression. For instance, *pleD* promotes flagellar disassembly by stimulating proteolytic turnover of the MS-ring protein FliF (77), but our results indicate that PleD is also stimulated through phosphorylation at D53 when flagellar assembly is disrupted (Fig 4A). Thus, cell cycle regulators that control flagellar assembly simultaneously act downstream of the flagellum in regulating holdfast production. This duality raises the intriguing possibility that specific environmental cues activate flagellar disassembly as part of a positive feedback loop that reinforces the commitment to differentiate into a stalked cell.

The developmental and mechanical pathways we identified each require a distinct diguanylate cyclase, suggesting that flagellar signaling converges to stimulate adhesion by modulating c-di-GMP levels (Fig 4C). Signaling through the mechanical pathway requires stator subunits that can productively engage with the rotor, and we propose that increased load on the flagellar filament induces conformational changes in the motor that activate the stator-associated diguanylate cyclase DgcB. This mechanism differs from contact-dependent activation of c-di-GMP synthesis by SadC in *Pseudomonas aeruginosa*, which requires disengagement of the MotCD stator(78). *P. aeruginosa* uses a MotAB stator system for swimming in liquid and a second, MotCD stator for swarming on surfaces(79). Thus, the signaling competency of engaged stators in *C. crescentus* could be an intrinsic feature of single-stator systems. Separately, c-di-GMP synthesis by PleD is part of a multi-tiered system controlling the master cell-cycle regulator CtrA in *C. crescentus*(53, 80), and we provided evidence that the status of flagellar assembly feeds into this developmental program by regulating PleD phosphorylation (Fig 5A). Whether the flagellum controls PleD phosphorylation through the DivJ-PleC kinase-phosphatase pair(57) or through a separate phosphorelay will require additional dissection of how *fss* genes link flagellar assembly to cell cycle progression.

Despite the apparent convergence of flagellar signaling on two diguanylate cyclases, *pleD*- and *dgcB* stimulate holdfast production by different mechanisms. When the mechanical pathway is bypassed by disrupting early stages of flagellar assembly, *shkA* and *pleD* are required for holdfast production, but *dgcB* is dispensable (Figs 2A and 3A). ShkA is a histidine kinase that is stimulated by c-di-GMP(81). It initiates a phosphotransfer that activates the transcriptional co-activator TacA, upregulating dozens of genes required for stalk biogenesis(47). A requirement for both *pleD* and *shkA* in the developmental pathway indicates that *pleD* controls holdfast production specifically through the *tacA* transcriptional program. In contrast, the *dgcB*-dependent c-di-GMP pool has been shown to act through direct activation of holdfast synthesis enzymes (40). Thus, PleD and DgcB likely act on different timescales, and we favor a model in which the sequential accumulation of c-di-GMP drives the transition to permanent attachment. In this scenario, increased load on the flagellar filament would activate DgcB, producing a transient burst of c-di-GMP that immediately stimulates holdfast production. Persistent filament obstruction would increase c-di-GMP levels sufficiently to destabilize the motor, leading to flagellar disassembly, activation of sustained c-di-GMP synthesis by PleD and the onset of stalked cell development (Fig 5C).

Using an unbiased screen to identify flagellar signaling genes has allowed us to propose a unified model for the mechanism by which the flagellum regulates holdfast production in *C. crescentus*. Intact flagellar motors respond to assembly defects in their associated filaments (Figs 2A and 5B), but perturbing the flagellum also influences the timing of holdfast production by altering cell cycle signaling (34)(Figs 2A and 5A). Candidate approaches specifically targeting developmental or tactile sensing phenomena have not accounted for presence of multiple pathways downstream of the flagellum. Though two pathways can be distinguished genetically in flagellar mutants, overlap between developmental, mechanical and other signaling pathways during actual surface encounters has likely confounded interpretations of how the flagellum regulates holdfast production. The complexity of these circuits underscores how bacterial behavior is not controlled by linear signaling pathways. Flagellar cues represent only a subset of the stimuli known to influence adhesion. Nutrient availability, redox homeostasis, chemotaxis and T4P dynamics all influence whether *C. crescentus* produces a holdfast. Disentangling how diverse signaling networks converge to regulate holdfast production has the power to illuminate how environmental information is integrated to control behavior.

## Materials and Methods

### Bacterial growth and genetic manipulation

Strains and plasmids used in this study are listed in Tables S4 and S5. Standard polymerase chain reaction (PCR) and Gibson assembly(82) methods were used for developing plasmid constructs. Strains, plasmids, primer sequences and details of construction are available upon request. *E. coli* cultures were grown in LB medium at 37°C supplemented with 1.5% (w/v) agar and 50 μg/mL kanamycin when necessary. Unless otherwise noted, *C. crescentus* cultures were grown at 30°C in PYE medium supplemented with 1.5% (w/v) agar, 3% (w/v) sucrose and 25 μg/mL kanamycin when necessary or in M2 defined medium supplemented with 0.15% (w/v) xylose (83). Plasmids were introduced into *C. crescentus* by electroporation. Unmarked deletions were constructed using *sacB*-based counterselection in sucrose as described previously(33).

### Genetic complementation of mutants

Mutants were complemented by genomic integration of the appropriate gene as a single copy at a neutral site (*xylX*) and under the gene’s native promoter. Specifically, predicted open reading frames were fused to their predicted promoter sequences and inserted into the NdeI/SacI site of pMT585 (pXGFPC-2)(84). Each promoter-gene cassette was inserted in reverse orientation to allow for transcription in the opposite direction relative to the *xylX* promoter upstream of the cloning site. Surface attachment and motility assays used to evaluate complementation are described below (Fig S4).

### Tn-Himar mutant library construction and mapping

Construction and mapping of the two barcoded transposon libraries was performed based on the procedure developed by Wetmore et al. as described previously(33, 85). Cells from 25 mL cultures of the APA_752 barcoded transposon pool that had been grown to mid-log phase in LB supplemented with kanamycin and 300 μM diaminopimelic acid (DAP) and 25mL of either the *C. crescentus* CB15 wild-type or Δ*flgH* strain that had been grown to mid-log phase in PYE were collected by centrifugation, washed twice with PYE containing 300 μM DAP, mixed and spotted together on a PYE agar plate containing 300 μM DAP. After incubating the plate overnight at room temperature, the cells were scraped from the plate, resuspended in PYE medium, spread onto 20, 150mm PYE agar plates containing kanamycin and incubated at 30°C for three days. Colonies from each plate were scraped into PYE medium and used to inoculate a 25mL PYE culture containing 5 μg/mL kanamycin. The culture was grown for three doublings, glycerol was added to 20%, and 1 mL aliquots were frozen at −80°C.

Library mapping was performed as described (85). Briefly, genomic DNA was isolated from three 1mL aliquots of each library. The DNA was sheared and ~300bp fragments were selected before end repair. A Y-adapter (Mod2_TS_Univ, Mod2_TruSeq) was ligated and used as a template for transposon junction amplification with the primers Nspacer_BarSeq_pHIMAR and either P7_mod_TS_index1 or P7_mod_TS_index2. 150-bp single end reads were collected on an Illumina HiSeq 2500 in rapid run mode, and the genomic insertion positions were mapped and correlated to a unique barcode using BLAT(86) and MapTnSeq.pl to generate a mapping file with DesignRandomPool.pl. All code is available at https://bitbucket.org/berkeleylab/feba/. Features of the barcoded transposon libraries can be found in Table S6.

### Adhesion profiling of barcoded Tn-Himar mutant libraries

Adhesion profiling was performed as in Hershey et al. (33) with slight modifications. Cells from 1mL aliquots of each barcoded transposon library were collected by centrifugation, resuspended in 1mL of M2X medium, and 300 μL was inoculated into a well of a 12-well microtiter plate containing 1.5mL M2X medium with 6-8 ~1 x 1 cm layers of cheesecloth. Microtiter plates containing selections were grown for 24hr at 30°C with shaking at 155rpm after which 150μL of the culture was passaged by inoculating into a well with 1.65mL fresh M2X containing cheesecloth. Cells from an additional 500μL of depleted medium were harvested by centrifugation and stored at −20°C for BarSeq analysis. Each passaging experiment was performed in triplicate, and passaging was performed sequentially for a total of five rounds of selection. Identical cultures grown in a plate without cheesecloth were used as a nonselective reference condition.

Cell pellets were used as PCR templates to amplify the barcodes in each sample using indexed primers(85). Amplified products were purified and pooled for multiplexed sequencing. 50bp single end reads were collected on an Illumina HiSeq4000. MultiCodes.pl, combineBarSeq.pl and FEBA.R were used to determine fitness by comparing the log2 ratios of barcode counts in each sample over the counts from a nonselective growth in M2X without cheesecloth. To evaluate mutant phenotypes in each screen, the replicates were used to calculate a mean fitness score for each gene after each passage. Mean fitness was summed across passages for each gene and ranked by either the lowest (WT background) or highest (Δ*flgH* background) summed fitness score.

### Surface attachment measurement by crystal violet (CV) staining

Overnight *C. crescentus* cultures grown in PYE were diluted to an OD_660_ of 0.5 with PYE, and 1.5 μL from each diluted starter culture was inoculated into the wells of a 48-well microtiter plate containing 450 μL M2X medium. The number of replicates for each experiment ranged from 5-8 and is indicated in the relevant figure legend. Plates were grown at 30°C with shaking at 155rpm for 17 hours after which the cultures were discarded, and the wells were washed thoroughly under a stream of tap water. Attached cells were stained by adding 500 μL of an aqueous solution containing 0.01% (w/v) crystal violet to each well and shaking the plates for 5 min. Excess dye was discarded, the wells were again washed under a stream of tap water and the remaining dye was dissolved by adding 500 μL of ethanol to each well. Staining was quantified by reading absorbance at 575nm using a Tecan Spark microplate reader. Each reading was corrected by subtracting the absorbance value for an uninoculated medium blank, the mean of the biological replicates for each strain was calculated and normalized to the mean value measured for the wild type background. To minimize day-to-day variation in the absolute CV staining values, each figure panel shows an internally controlled experiment with all measurements taken from the same plate on the same day. The collection of strains shown in each figure was assayed together on at least four independent days and a representative dataset is shown.

### Holdfast staining with fluorescent wheat germ agglutinin (fWGA)

2mL of M2X medium was inoculated to achieve a starting OD_660_ of 0.001 using saturated starter cultures grown in PYE. After growing to an OD_660_ of 0.07-0.1, 400 μL of each culture was added to a fresh 1.5mL ependorf tube containing 1 μL of a 2mg/mL solution of WGA conjugated with Alexa594. After a 5min incubation at room temperature in the dark, cells were harvested at 6k x g, washed with distilled water, and resuspended in the residual liquid after centrifugation. 1 μL was spotted onto a glass slide and a cover slip was placed on top. Imaging was performed with a Leica DM5000 microscope equipped with an HCX PL APO63X/1.4-numerical-aperture Ph3 objective. A red fluorescent protein filter (Chroma set 41043) was used to visualize WGA foci. Cells with a holdfast were counted manually on five separate days. A minimum of 95 cells were counted for each biological replicate.

For direct staining of liquid cultures without centrifugation, cells were grown as described above. 400 μL was removed and imaged using the standard protocol described above, and 0.8 μL from a 2mg/mL fWGA solution was added directly to the remaining culture. Cultures were shaken with the dye for 5 min in the dark after which 1.5 μL was spotted on a microscope slide, covered and imaged immediately as described above.

### Analysis of holdfast polysaccharide in spent medium

Holdfast release was analyzed as described previously(43). 10mL cultures grown for 24 hours in M2X were centrifuged at 7k x g for 10 min. 2mL of supernatant (spent medium) was moved to a fresh tube, 3mL of 100% ethanol was added, and the mixture was incubated overnight at 4°C. Precipitate was isolated by centrifugation for 1hr at 18,000 x g and suspended in 50 μL TU buffer (10mM Tris-HCl pH 8.2, 4M urea). 2-fold serial dilutions were prepared, and 3 μL of each dilution was spotted on nitrocellulose to absorb for 20 min. The membrane was then blocked overnight with 5% bovine serum albumin (BSA) dissolved in TBST buffer (20mM Tris-HCl pH 8.0, 137mM NaCl, 2.7mM KCl, 0.1% Tween 20) followed by a 1hr incubation with 5% BSA in TBST containing 1.5 ug/mL fWGA. The membrane was washed with TBST and imaged with a BioRad ChemiDoc imager using the Alexa Fluor 647 setting.

### Soft-agar motility assay

Overnight cultures grown in PYE were diluted to OD_660_ = 0.5, and 1.5 μL was pipetted into a PYE plate containing 0.3% agar. Plates were sealed with parafilm, incubated for 72 hours at 30°C and photographed.

### Flagellar filament staining

2mL of PYE medium was inoculated to a starting OD_660_ = 0.05 using saturated overnight starter cultures and grown at 30°C to mid-log phase (OD_660_ = 0.3 – 0.4). 500 μL of each culture was mixed with 0.5 μL of a 2mg/mL solution of Alexa488-maleimide in DMSO and incubated for 10min in the dark. Cells were harvested by centrifuging at 6k x g for 1.5 min, washed with 500 μL PYE, re-centrifuged and suspended in 500 μL PYE. 1 μL of the stained cell suspension was spotted on a pad of PYE solidified with 1% agarose. Imaging was performed as above but with the use of green fluorescent protein filter (Chroma set 41017) for flagellin visualization. Note that Alexa-488 maleimide cross-reacts with the holdfast.

### Microscopic analysis of swimming behavior

2mL of PYE was inoculated to OD_660_ = 0.1 with saturated overnight starters and grown at 30°C to OD_660_ = 0.4 – 0.5. A 2.69% (w/v) solution of 2.0 μm polystyrene spacer beads (Polysciences) was diluted 1000-fold in 1mL PYE. Equal volumes of liquid culture and diluted spacer beads were mixed and spotted on a slide. Dark field images were collected at 100ms intervals for 30s using a Leica 40X PH2 objective. Maximum projections from each time series are presented.

### Mapping of suppressing mutations in ΔfssA

To isolate motile suppressors, 1.5 μL from a saturated Δ*fssA* culture grown in PYE was spotted into PYE plates containing 0.3% agar. Plates were sealed and incubated for 96 hours at 30°C. Cells from the leading edge of spreading flares (Fig 4A) were streaked onto a standard PYE plate, and the plates were incubated at 30°C for 72 hours. A single colony was inoculated into PYE broth and grown to saturation. To avoid isolating siblings, only one suppressor was isolated from each initial soft-agar spotting. Genomic DNA from fourteen suppressors as well as the original Δ*fssA* parent background was isolated as described above. Libraries were prepared based on the Illumina Nextera protocol and single end reads were collected using the NextSeq 550 platform at the Microbial Genome Sequencing Center (MiGS, Pittsburgh, USA). Mutations were identified using breseq(87) with the *C. crescentus* NA1000 as a reference genome (GenBank CP001340).

### Homology modelling of CcMotAB

To develop a structural model of the *C. crescentus stator*, we used the MotA and MotB protein sequences (accessions CCNA_00787 & CCNA_01644) to search a protein structure fold library using the HHpred/HHSearch package for homology detection (88) within the Phyre2 pipeline(89). For *C. crescentus* MotB, this approach yielded high confidence (>99%) structural models of the N- and C-terminal halves of the protein that aligned to multiple published MotB structures, including the N-terminus of *C. jejuni* MotB (PDB ID: 6YKM). The *C. crescentus MotA* model aligned to the entire length of *Campylobacter jejuni* MotA (PDB ID: 6YKM) with high confidence (100%). The coordinates of the *Cc*MotA and N-terminal *Cc*MotB homology models were used to build a 5:2 MotA:MotB complex by aligning to the 6YKM *C. jejuni* MotAB model.

### β-galactosidase assay

Strains carrying a P_*hfiA*_-*lacZ* transcriptional reporter (35) were inoculated from colonies on PYE-agar plates into 2 ml M2X medium and grown shaking at 200 RPM overnight at 30°C. Overnight cultures were diluted in 2 ml of fresh M2X to an OD_660_ of 0.05 and grown for approximately 6 hours to early exponential phase. These cultures were then diluted again into 2 ml fresh M2X to an OD_660_ of 0.001 and grown for 17 hours to a final OD660nm of 0.1. *β*-galactosidase activity was then was measured colorimetrically as previously described(35). Briefly, 0.15 ml of cells were permeabilized by vortexing with 50 ul of chloroform and 50 ul of PYE broth as an emulsifier. 600 ul of Z-buffer (60 mM Na_2_HPO_4_, 40 mM NaH_2_PO_4_, 10 mM KCl, 1 mM MgSO_4_) was added, followed by 200 ul ONPG in 0.1 M KPO_4_. After 5 minutes at room temperature, reactions were quenched with 1 ml of 1 M Na_2_CO_3_ and absorbance at 420nm was used to calculate *β*-galactosidase activity.

### Data availability

Sequence data have been deposited in the NCBI Sequence Read Archive (SRA) with the following project accessions. For wild-type *C. crescentus*, PRJNA640825 contains the sequence data used to map the barcoded TnHimar library, and PRJNA640525 contains barcoded amplicon sequences collected after passaging in cheesecloth. For the Δ*flgH* mutant, PRJNA640725 contains the sequencing data used to map the barcoded TnHimar library, and PRJNA641033 contains barcoded amplicon sequences collected after passaging in cheesecloth. PRJNA672134 contains whole genome sequencing data for the Δ*fssA* parent strain and 14 motile suppressors.

## Acknowledgements

We thank members of the Crosson laboratory, Howard Shuman and Phoebe Rice for helpful discussions. We thank Miette Hennessey for assistance in constructing the *lacZ* reporter strains. This work was supported by NIH grant R01GM087353 to S.C. D.M.H. is supported by the Helen Hay Whitney Foundation.

## Supplemental materials

***Table S1** Mutated genes that produce a hyper-adhesive phenotype during adhesion profiling* Mean fitness scores for each successive passage (P1-P5) of the barcoded CB15 Tn-Himar mutant library in cheesecloth are shown along with a nonselective control (P0). Negative values indicate hyper-adhesive strains that are depleted more rapidly than the bulk population.

***Table S2** Mutated genes that suppress hyper-adhesion in the* ΔflgH *background during adhesion profiling* Mean fitness scores for each successive passage (P1-P5) of the barcoded Δ*flgH* Tn-Himar mutant library in cheesecloth are shown along with a nonselective control (P0). Positive values indicate strains with reduced adhesion that are depleted less rapidly than the bulk population.

***Table S3** Statistical analysis of strain comparisons shown in Figure 2*

The results of ANOVA followed by Tukey’s multiple comparison test for 11 critical strains in Fig 2 is shown. Adjusted P values for each comparison are highlighted in blue, and statistical significance is highlighted in green. (*) – P < 0.05; (**) – P < 0.01; (***) – P < 0.001; (****) – P < 0.0001.

***Table S4** Plasmids used in this study*

***Table S5** Strains used in this study*

***Table S6** Features of the barcoded transposon libraries used in this study*

***Figure S1** Surface contact independent activation of holdfast production in* ΔflgH A) Shedding of holdfast polysaccharide into spent medium. Δ*flgH* releases a holdfast specific fWGA reactive material into the spent medium during growth in M2X liquid. B) Comparison showing the fraction of holdfast producing cells in wild type and Δ*flgH* backgrounds. Centrifugation has no effect on holdfast production in either strain. C) Micrographs of wild type and Δ*flgH* mutant cells taken immediately after direct staining of holdfast in liquid cultures.

***Figure S2** Confirmation of the two-pathway model for flagellar control of holdfast production* A) CV staining of mutants from Fig 2D grown in PYE. The loss of adhesion when the mechanical (Δ*motB*) and developmental (Δ*pleD*) pathways are inactivated simultaneously is not specific to defined M2X medium. Mean values from six biological replicates are shown along with their associated standard deviations. B) CV stain confirming the placement of *motA* in the mechanical pathway. Δ*motA* reduces hyper-adhesion specifically in late flagellar mutants, mirroring the Δ*motB* suppression pattern. Mean values from seven biological replicates are shown along with their associated standard deviations. C) CV stain showing additive effects of disrupting *shkA* and *motB* simultaneously. The results confirm the placement of *shkA* and *motB* in separate signaling pathways. Mean values from six biological replicates are shown along with their associated standard deviations. D) CV stain showing additive effects of disrupting *shkA* and *dgcB* simultaneously. The results confirm the placement of *shkA* and *dgcB* in separate signaling pathways. Mean values from six biological replicates are shown along with their associated standard deviations.

***Figure S3** Additional analysis of the* ΔfssA *phenotype* A) Mapping of mutations that suppress the Δ*fssA* motility phenotype by whole genome sequencing. The suppressor numbers correspond to files in SRA accession PRJNA672134. Nucleotide positions correspond to coordinates in the NA1000 genome. B) Soft-agar motility assay showing that the Δ*fssB* motility defect is epistatic to the Δ*fssA* phenotype. Motile suppressors do not appear in the Δ*fssA* Δ*fssB* double mutant after a 96 hour incubation. C) CV staining experiment showing that the Δ*fssA* and Δ*fssB* are not additive for suppression of hyper-adhesion in the Δ*flgH* mutant. Mean values from six biological replicates are shown along with their associated standard deviations.

***Figure S4** Genetic complementation of surface attachment and motility phenotypes* A) Complementation of flagellar hierarchy mutants in soft agar. B) Complementation of hyper-adhesion in flagellar mutants. C) Complementation of crystal violet staining for *fss* mutants in Δ*flgH* background. D, E) Restoration of motility phenotypes in soft agar for *fss* mutants.

## Notes

### Competing Interest Statement

The authors have declared no competing interest.

### Summary of Updates

This version of the manuscript has been revised in response to external peer review.

