## Supplemental Materials for "Flagellar perturbations activate adhesion through two distinct pathways in *Caulobacter crescentus*"

***Table S1*** *Mutated genes that produce a hyper-adhesive phenotype during adhesion profiling* Mean fitness scores for each successive passage (P1-P5) of the barcoded CB15 Tn-Himar mutant library in cheesecloth are shown along with a nonselective control (P0). Negative values indicate hyper-adhesive strains that are depleted more rapidly than the bulk population.

| Gene | P0 | P1 | P2 | P3 | P4 | P5 |
|---|---|---|---|---|---|---|
| CCNA_03484; NfeD-related membrane protease regulator | -0.18 | -2.52 | -3.62 | -5.07 | -6.14 | -6.34 |
| CCNA_02666; chemotactic signal-response protein CheL | 0.22 | -2.30 | -3.76 | -4.50 | -4.42 | -4.49 |
| CCNA_01526; flagellar biosynthesis regulatory protein FlaF | -0.12 | -1.13 | -3.55 | -4.40 | -4.39 | -5.25 |
| CCNA_00747; cyclic-di-GMP phosphodiesterase, flagellum assembly factor TipF | -0.12 | -1.76 | -3.34 | -4.12 | -4.49 | -4.32 |
| CCNA_02142; flagellar basal-body rod protein FlgF | -0.05 | -1.41 | -2.88 | -4.41 | -4.64 | -4.59 |
| CCNA_00050; apolipoprotein N-acyltransferase Lnt | 0.17 | -1.68 | -2.06 | -3.61 | -4.31 | -4.43 |
| CCNA_03538; NAD-dependent epimerase/ dehydratase | 0.01 | -2.80 | -3.01 | -3.27 | -3.59 | -3.07 |
| CCNA_00952; flagellar assembly protein FlbE/FliH | 0.07 | -1.72 | -3.81 | -3.99 | -3.46 | -2.72 |
| CCNA_02664; flagellar assembly regulator FliX | -0.09 | -1.22 | -3.40 | -3.22 | -3.87 | -3.51 |
| CCNA_02946; spsF-related cytidylyltransferase | -0.11 | -1.53 | -3.21 | -3.29 | -3.36 | -3.45 |
| CCNA_01002; flagellar biosynthesis protein FliP | 0.02 | -2.16 | -3.03 | -3.43 | -3.58 | -2.61 |
| CCNA_01005; flagellar basal-body rod protein FlgC | -0.22 | -1.42 | -2.68 | -3.39 | -3.78 | -3.44 |
| CCNA_01004; flagellar basal-body rod protein FlgB | 0.20 | -1.32 | -2.22 | -3.67 | -3.91 | -3.19 |
| CCNA_00954; AAA-family response regulator FlbD | 0.15 | -1.75 | -2.74 | -2.82 | -3.45 | -3.43 |
| CCNA_02411; soluble lytic transglycosylase PleA | -0.01 | -1.54 | -2.84 | -3.04 | -3.19 | -3.37 |
| CCNA_02667; flagellar basal-body protein FlbY | 0.00 | -1.24 | -2.55 | -3.44 | -3.24 | -3.35 |
| CCNA_02140; flagellar motor switch protein FliM | 0.15 | -1.98 | -2.93 | -3.10 | -2.77 | -3.05 |
| CCNA_00943; flagellar hook-associated protein FlaN | -0.01 | -1.13 | -2.69 | -3.23 | -3.28 | -3.50 |
| CCNA_02860; DnaJ-class molecular chaperone | -0.28 | -0.77 | -1.93 | -3.44 | -3.43 | -4.15 |
| CCNA_00821; hypothetical protein | 0.04 | -2.05 | -2.30 | -2.36 | -3.33 | -3.48 |
| CCNA_00956; flagellar biosynthesis protein FlhA | 0.04 | -1.71 | -2.75 | -3.14 | -2.83 | -2.96 |
| CCNA_02855; PepSY peptidase propeptide domain protein | -0.02 | -2.09 | -1.22 | -2.37 | -3.64 | -4.02 |
| CCNA_02143; flagellar basal-body rod protein FlgG | 0.04 | -1.15 | -2.54 | -3.07 | -3.20 | -3.22 |
| CCNA_02144; flagella basal body P ring formation protein FlgA | -0.02 | -1.26 | -2.12 | -3.41 | -3.45 | -2.91 |
| CCNA_02315; Cro/CI-family transcriptional regulator | -0.04 | -0.87 | -2.13 | -2.65 | -3.54 | -3.72 |
| CCNA_00946; basal-body rod modification protein FlgD | 0.23 | -0.57 | -1.81 | -2.57 | -3.42 | -4.48 |
| CCNA_00953; flagellar motor switch protein FliN | 0.09 | -1.59 | -2.27 | -2.93 | -3.02 | -2.82 |
| CCNA_01532; regulatory protein FlaY | 0.01 | -1.02 | -2.24 | -2.95 | -3.20 | -3.17 |
| CCNA_00947; flagellar hook protein FlgE | 0.02 | -0.99 | -2.13 | -2.95 | -3.19 | -3.14 |
| CCNA_00446; chemotaxis receiver domain protein CheYII | 0.02 | -1.09 | -1.92 | -2.71 | -2.97 | -3.67 |
| CCNA_00942; flagellar hook-associated protein FlgL | 0.11 | -1.15 | -2.22 | -2.72 | -3.01 | -3.15 |
| CCNA_01003; flagellar biosynthesis protein FliO | 0.11 | -1.13 | -2.67 | -2.54 | -3.30 | -2.58 |
| CCNA_01131; flagellar biosynthesis protein FlhB | -0.13 | -1.98 | -2.05 | -2.80 | -2.57 | -2.67 |
| CCNA_00668; capsular polysaccharide biosynthesis protein | -0.10 | -1.21 | -1.47 | -2.73 | -3.51 | -3.13 |

| | | | | | | |
|---|---|---|---|---|---|---|
| CCNA_01445; hypothetical protein | -0.16 | -1.72 | -2.12 | -2.25 | -2.85 | -2.88 |
| CCNA_02949; S-adenosylmethionine-dependent methyltransferase related protein | -0.06 | -1.20 | -2.16 | -2.82 | -2.70 | -2.87 |
| CCNA_02463; UDP glucuronic acid epimerase | -0.12 | -2.20 | -1.98 | -2.47 | -2.18 | -2.70 |
| CCNA_00944; flagellar hook length determination protein | -0.23 | -2.05 | -1.05 | -2.84 | -3.11 | -2.47 |
| CCNA_00459; Na+/H+ antiporter NhaA | -0.01 | -0.88 | -1.80 | -2.64 | -3.09 | -3.02 |
| CCNA_02950; protoporphyrinogen oxidase-related protein | 0.07 | -0.87 | -1.89 | -2.38 | -3.09 | -3.18 |
| CCNA_01524; flagellar modification protein FlbA | 0.02 | -1.02 | -2.16 | -2.48 | -2.91 | -2.76 |
| CCNA_02951; WbqC-like family protein | 0.09 | -0.51 | -1.87 | -2.52 | -3.27 | -3.16 |
| CCNA_00823; LuxR-like DNA-binding protein | -0.01 | -1.37 | -1.44 | -2.19 | -3.26 | -2.91 |
| CCNA_01219; putative cytosolic protein | -0.54 | -0.43 | -1.66 | -2.16 | -3.33 | -3.59 |
| CCNA_00950; flagellar M-ring protein FliF | 0.12 | -1.42 | -2.18 | -2.02 | -2.91 | -2.60 |
| CCNA_02145; flagellar L-ring protein FlgH | 0.07 | -0.89 | -2.04 | -2.70 | -2.86 | -2.61 |
| CCNA_03135; flagellar protein export ATPase FliL | 0.05 | -1.39 | -2.03 | -2.55 | -2.65 | -2.15 |
| CCNA_00233; UDP-N-acetylglucosamine 4,6-dehydratase | 0.23 | -0.98 | -2.33 | -2.43 | -2.30 | -2.71 |
| CCNA_03136; flagellar biosynthesis chaperone FliJ | 0.02 | -1.37 | -2.01 | -2.45 | -2.32 | -2.51 |
| CCNA_01198; dTDP-4-dehydrorhamnose reductase | -0.04 | -0.88 | -1.89 | -2.65 | -2.55 | -2.67 |
| CCNA_00444; chemotaxis protein methyltransferase CheRI | 0.07 | -0.87 | -1.67 | -2.29 | -2.71 | -3.01 |
| CCNA_02947; spsG-related polysaccharide biosynthesis protein | 0.11 | -0.94 | -2.26 | -2.41 | -2.52 | -2.37 |
| CCNA_02241; YjgF/YER057c/UK114-family protein | -0.04 | -1.26 | -2.11 | -2.57 | -2.12 | -2.41 |
| CCNA_00501; glucosyltransferase | -0.04 | -1.63 | -2.09 | -2.31 | -2.06 | -2.26 |
| CCNA_03265; CBS pair-family sensor histidine kinase/receiver domain protein | -0.03 | -1.32 | -2.21 | -2.01 | -2.04 | -2.54 |
| CCNA_00234; WecE-family cell wall biogenesis enzyme | 0.07 | -0.63 | -1.89 | -2.45 | -2.42 | -2.56 |
| CCNA_02138; flagellar motility protein MotE | -0.05 | -1.26 | -2.04 | -1.86 | -2.43 | -2.17 |
| CCNA_01130; flagellar biosynthesis protein FliR | -0.08 | -0.90 | -1.40 | -2.31 | -2.59 | -2.49 |
| CCNA_00447; chemotaxis protein CheD | 0.19 | -0.66 | -1.30 | -2.57 | -2.48 | -2.63 |
| CCNA_00390; ADP-heptose--LPS heptosyltransferase | 0.10 | -1.61 | -1.93 | -1.80 | -2.30 | -1.97 |
| CCNA_02127; bifunctional lysylphosphatidylglycerol flippase/synthetase MprF | 0.16 | -1.01 | -0.61 | -2.22 | -2.69 | -2.77 |
| CCNA_01675; outer membrane translocation and assembly protein TamA | 0.02 | -1.40 | -1.99 | -2.03 | -1.83 | -1.97 |
| CCNA_01446; LL-diaminopimelate aminotransferase | 0.09 | -1.05 | -1.65 | -2.22 | -2.16 | -2.10 |
| CCNA_01058; helix-turn-helix transcriptional regulator | -0.03 | -0.65 | -1.16 | -1.50 | -2.86 | -2.94 |
| CCNA_02282; VanZ family protein | 0.03 | -0.63 | -1.25 | -1.57 | -2.32 | -3.16 |
| CCNA_02961; NeuB-family N-acetylneuraminate synthase | 0.07 | -1.00 | -1.69 | -2.04 | -2.09 | -1.98 |
| CCNA_01430; hypothetical protein | 0.05 | -2.04 | -1.31 | -2.04 | -1.82 | -1.59 |
| CCNA_03577; DNA polymerase I | 0.02 | -0.46 | -0.70 | -1.74 | -2.76 | -2.82 |
| CCNA_03289; type II toxin-antitoxin system HicB family antitoxin | -0.17 | -0.19 | -0.35 | -1.76 | -3.19 | -2.97 |
| CCNA_03507; GGDEF/EAL phosphodiesterase PdeA | 0.00 | -1.23 | -1.69 | -1.57 | -1.88 | -1.85 |
| CCNA_02665; flagellar P-ring protein FlgI | 0.12 | -0.67 | -1.47 | -2.05 | -2.10 | -1.93 |
| CCNA_01373; hypothetical protein | 0.06 | -0.20 | -0.37 | -1.39 | -3.00 | -3.26 |

| | | | | | | |
|---|---|---|---|---|---|---|
| CCNA_01676; autotransporter translocation and assembly factor TamB | -0.01 | -1.24 | -1.60 | -1.59 | -1.90 | -1.84 |
| CCNA_02525; aspartate carbamoyltransferase | -0.29 | 0.17 | -1.75 | -1.56 | -1.89 | -3.13 |
| CCNA_02881; oxoglutarate semialdehyde dehydrogenase | 0.11 | -0.97 | -1.38 | -2.04 | -1.73 | -2.03 |

***Table S2*** *Mutated genes that suppress hyper-adhesion in the Δ*flgH *background during adhesion profiling* Mean fitness scores for each successive passage (P1-P5) of the barcoded Δ*flgH* Tn-Himar mutant library in cheesecloth are shown along with a nonselective control (P0). Positive values indicate strains with reduced adhesion that are depleted less rapidly than the bulk population.

| Gene | P0 | P1 | P2 | P3 | P4 | P5 |
|---|---|---|---|---|---|---|
| CCNA_02513; holdfast synthesis protein HfsA | -0.11 | 1.27 | 3.36 | 5.43 | 7.46 | 9.03 |
| CCNA_02512; polysaccharide autokinase-related protein HfsB | 0.00 | 1.29 | 3.35 | 5.35 | 7.19 | 8.63 |
| CCNA_02514; polysaccharide secretin protein HfsD | -0.06 | 1.20 | 3.19 | 5.23 | 7.22 | 8.79 |
| CCNA_02567; sensory transduction histidine kinase PleC | -0.04 | 0.38 | 3.08 | 5.34 | 7.51 | 9.29 |
| CCNA_00094; WecG/TagA-family glycosyltransferase HfsJ | 0.02 | 1.29 | 3.26 | 5.19 | 7.10 | 8.65 |
| CCNA_02360; holdfast glycosyl transferase family 2 protein HfsL | -0.14 | 1.15 | 2.99 | 5.14 | 7.11 | 8.50 |
| CCNA_02509; glycosyltransferase HfsG | 0.00 | 1.07 | 3.02 | 5.10 | 7.01 | 8.36 |
| CCNA_02510; oligosaccharide deacetylase HfsH | -0.16 | 1.27 | 3.04 | 4.86 | 6.75 | 8.12 |
| CCNA_03043; type IV pilin protein PilA | 0.12 | 0.95 | 2.82 | 4.58 | 6.15 | 7.16 |
| CCNA_01926; GGDEF diguanylate cyclase DgcB | 0.16 | 0.99 | 2.73 | 4.61 | 6.13 | 7.17 |
| CCNA_00137; hybrid two-component histidine kinase/receiver protein ShkA | -0.19 | 1.16 | 2.74 | 4.21 | 5.59 | 6.64 |
| CCNA_01644; chemotaxis protein MotB | -0.01 | 1.35 | 2.61 | 4.18 | 5.40 | 6.27 |
| CCNA_02137; TPR-containing protein FssB | 0.05 | 1.03 | 2.30 | 3.86 | 5.18 | 6.05 |
| CCNA_01117; conserved hypothetical protein FssA | 0.25 | 0.94 | 2.38 | 3.83 | 5.07 | 5.94 |
| CCNA_00948; CtrA inhibitory protein SciP | 0.43 | 1.27 | 2.21 | 3.83 | 4.79 | 5.57 |
| CCNA_02141; flagellar protein FliL | 0.03 | 0.89 | 2.14 | 3.56 | 4.92 | 5.79 |
| CCNA_02298; zinc-finger pilus regulatory protein ZitP | 0.08 | 1.15 | 2.75 | 3.75 | 4.64 | 5.01 |
| CCNA_03424; AAA-family response regulator TacA | -0.03 | 0.73 | 1.93 | 3.42 | 4.95 | 5.89 |
| CCNA_02125; polar development protein PodJ | -0.07 | 0.43 | 1.62 | 3.08 | 4.80 | 6.57 |
| CCNA_02546; GGDEF/response regulator protein PleD | -0.08 | 0.60 | 2.02 | 3.32 | 4.48 | 5.31 |
| CCNA_02257; fliN family protein FliN2 | 0.29 | 1.15 | 2.18 | 3.41 | 4.21 | 4.70 |
| CCNA_02087; deoxyguanosinetriphosphate triphosphohydrolase | 0.13 | 0.70 | 2.10 | 3.36 | 4.40 | 5.01 |
| CCNA_02712; holdfast attachment protein HfaB | -0.22 | 0.80 | 1.90 | 3.10 | 4.34 | 5.28 |
| CCNA_01171; histidine phosphotransferase ShpA | -0.37 | 0.92 | 1.56 | 2.78 | 4.15 | 4.97 |
| CCNA_03713; RNA polymerase sigma-54 factor RpoN | 0.03 | -0.03 | 1.80 | 2.92 | 4.02 | 4.73 |
| CCNA_03803; GNAT-family holdfast biogenesis protein HfsK | 0.04 | 0.52 | 1.60 | 2.72 | 3.87 | 4.63 |
| CCNA_02711; holdfast attachment protein HfaA | 0.09 | 0.65 | 1.56 | 2.47 | 3.72 | 4.38 |
| CCNA_02332; chemotaxis receiver protein CheYIII | -0.02 | 0.47 | 1.39 | 2.79 | 3.67 | 4.43 |
| CCNA_00787; chemotaxis protein MotA | -0.06 | 0.49 | 1.13 | 2.40 | 3.61 | 4.41 |
| CCNA_02508; oligosaccharide translocase/flippase HfsF | -0.09 | 0.70 | 1.51 | 2.49 | 3.37 | 3.87 |
| CCNA_03830; HlyD family secretion protein | -0.11 | 0.83 | 1.36 | 2.57 | 3.28 | 3.57 |
| CCNA_00446; chemotaxis receiver domain protein CheYII | 0.02 | 1.06 | 1.21 | 2.20 | 3.08 | 3.77 |
| CCNA_03831; major facilitator superfamily transporter | 0.02 | 0.69 | 1.17 | 2.36 | 3.01 | 3.41 |
| CCNA_00676; PSK-family transcription factor | -0.58 | 0.18 | 0.86 | 1.80 | 3.19 | 4.51 |

| | | | | | | |
|---|---|---|---|---|---|---|
| CCNA_01379; type I secretion outer membrane protein RsaFb | -0.13 | 0.91 | 1.13 | 2.19 | 2.93 | 3.26 |
| CCNA_03585; chemotaxis receiver domain protein CheYIV | -0.70 | 0.90 | 1.24 | 2.14 | 2.86 | 3.16 |
| CCNA_03586; chemotaxis protein methyltransferase CheRIII | 0.20 | 0.54 | 1.26 | 2.02 | 2.74 | 3.18 |
| CCNA_01280; cell development regulator SpmY | -0.10 | 0.05 | 1.44 | 2.01 | 2.57 | 3.12 |
| CCNA_01226; OstA family protein | -0.04 | 0.55 | 1.56 | 1.99 | 2.27 | 2.40 |
| CCNA_01213; YjgP/YjgQ family membrane permease | -0.15 | 0.24 | 0.99 | 2.21 | 2.59 | 2.73 |
| CCNA_03042; pilus assembly prepilin peptidase CpaA | 0.03 | 0.33 | 1.12 | 1.78 | 2.55 | 2.98 |
| CCNA_03525; hypothetical protein | -1.60 | 1.38 | 1.26 | 1.58 | 1.77 | 2.21 |
| CCNA_03395; two-component receiver domain protein | -0.17 | 0.59 | 1.10 | 1.71 | 2.31 | 2.46 |
| CCNA_02255; lysozyme-family localization factor SpmX | -0.50 | 0.10 | 0.87 | 1.73 | 2.57 | 2.86 |
| CCNA_02358; flavin reductase family protein | -0.28 | 0.41 | 0.46 | 1.15 | 2.33 | 3.77 |
| CCNA_00193; 3-isopropylmalate dehydrogenase | -0.17 | 0.24 | 0.40 | 1.09 | 2.20 | 3.42 |
| CCNA_02409; hybrid sensor histidine kinase/receiver protein | 0.17 | 0.39 | 0.82 | 1.54 | 1.99 | 2.21 |
| CCNA_03552; polysaccharide deacetylase CpaM | -0.24 | 0.48 | 0.94 | 1.44 | 1.97 | 2.08 |
| CCNA_02866; YhaH-family conserved membrane protein | 0.11 | -0.68 | 0.26 | 1.08 | 2.40 | 3.80 |
| CCNA_00674; NAD(FAD)-utilizing dehydrogenase | -0.40 | 0.11 | 0.31 | 0.88 | 1.93 | 3.22 |

***Table S3*** *Statistical analysis of strain comparisons shown in Figure 2*

The results of ANOVA followed by Tukey's multiple comparison test for 11 critical strains in Fig 2 is shown. Adjusted P values for each comparison are highlighted in blue, and statistical significance is highlighted in green. (*) – P < 0.05; (**) – P < 0.01; (***) – P < 0.001; (****) – P < 0.0001.

| **Crystal violet staining (Figs 2A and 2B)** | | | | | | | | | | | |
|---|---|---|---|---|---|---|---|---|---|---|---|
| | **WT** | **Δ*pleD*** | **Δ*motB*** | **Δ*pleD* Δ*motB*** | **Δ*fliF*** | **Δ*fliF* Δ*pleD*** | **Δ*fliF* Δ*motB*** | **Δ*flgH*** | **Δ*flgH* Δ*pleD*** | **Δ*flgH* Δ*motB*** | **Δ*flgH* Δ*pleD* Δ*motB*** |
| **WT** | X | ns | *** | **** | **** | **** | **** | **** | ns | ns | **** |
| **Δ*pleD*** | 0.9635 | X | **** | **** | **** | **** | **** | **** | ns | ns | **** |
| **Δ*motB*** | 0.0006 | <0.0001 | X | **** | **** | **** | **** | **** | ** | * | **** |
| **Δ*pleD* Δ*motB*** | <0.0001 | <0.0001 | <0.0001 | X | **** | ns | **** | **** | **** | **** | ns |
| **Δ*fliF*** | <0.0001 | <0.0001 | <0.0001 | <0.0001 | X | **** | ns | **** | **** | **** | **** |
| **Δ*fliF* Δ*pleD*** | <0.0001 | <0.0001 | <0.0001 | >0.9999 | <0.0001 | X | **** | **** | **** | **** | ns |
| **Δ*fliF* Δ*motB*** | <0.0001 | <0.0001 | <0.0001 | <0.0001 | 0.8699 | <0.0001 | X | **** | **** | **** | **** |
| **Δ*flgH*** | <0.0001 | <0.0001 | <0.0001 | <0.0001 | <0.0001 | <0.0001 | <0.0001 | X | **** | **** | **** |
| **Δ*flgH* Δ*pleD*** | 0.9992 | 0.5788 | 0.0073 | <0.0001 | <0.0001 | <0.0001 | <0.0001 | <0.0001 | X | ns | **** |
| **Δ*flgH* Δ*motB*** | 0.9889 | 0.3786 | 0.018 | <0.0001 | <0.0001 | <0.0001 | <0.0001 | <0.0001 | >0.9999 | X | **** |
| **Δ*flgH* Δ*pleD* Δ*motB*** | <0.0001 | <0.0001 | <0.0001 | >0.9999 | <0.0001 | >0.9999 | <0.0001 | <0.0001 | <0.0001 | <0.0001 | X |

| **Holdfast count (Fig 2C)** | | | | | | | | | | | |
|---|---|---|---|---|---|---|---|---|---|---|---|
| | **WT** | **Δ*pleD*** | **Δ*motB*** | **Δ*pleD* Δ*motB*** | **Δ*fliF*** | **Δ*fliF* Δ*pleD*** | **Δ*fliF* Δ*motB*** | **Δ*flgH*** | **Δ*flgH* Δ*pleD*** | **Δ*flgH* Δ*motB*** | **Δ*flgH* Δ*pleD* Δ*motB*** |
| **WT** | X | ns | **** | ** | **** | * | **** | **** | * | *** | ** |
| **Δ*pleD*** | >0.9999 | X | **** | * | **** | * | **** | **** | ** | **** | * |
| **Δ*motB*** | <0.0001 | <0.0001 | X | **** | **** | **** | **** | *** | ns | ns | **** |
| **Δ*pleD* Δ*motB*** | 0.005 | 0.0153 | <0.0001 | X | **** | ns | **** | **** | **** | **** | ns |
| **Δ*fliF*** | <0.0001 | <0.0001 | <0.0001 | <0.0001 | X | **** | ns | **** | **** | **** | **** |
| **Δ*fliF* Δ*pleD*** | 0.0133 | 0.0382 | <0.0001 | >0.9999 | <0.0001 | X | **** | **** | **** | **** | ns |

| | | | | | | | | | | | |
|---|---|---|---|---|---|---|---|---|---|---|---|
| ***ΔfliF<br>ΔmotB*** | <0.0001 | <0.0001 | <0.0001 | <0.0001 | 0.666 | <0.0001 | X | **** | **** | **** | **** |
| ***ΔflgH*** | <0.0001 | <0.0001 | 0.0001 | <0.0001 | <0.0001 | <0.0001 | <0.0001 | X | **** | **** | **** |
| ***ΔflgH<br>ΔpleD*** | 0.0181 | 0.0059 | 0.0786 | <0.0001 | <0.0001 | <0.0001 | <0.0001 | <0.0001 | X | ns | **** |
| ***ΔflgH<br>ΔmotB*** | 0.0001 | <0.0001 | 0.8579 | <0.0001 | <0.0001 | <0.0001 | <0.0001 | <0.0001 | 0.8857 | X | **** |
| ***ΔflgH<br>ΔpleD<br>ΔmotB*** | 0.0059 | 0.018 | <0.0001 | >0.9999 | <0.0001 | >0.9999 | <0.0001 | <0.0001 | <0.0001 | <0.0001 | X |

### *P*$_{hfiA}$-*lacZ* reporter activity (Fig 2D)

| | **WT** | ***ΔpleD*** | ***ΔmotB*** | ***ΔpleD<br>ΔmotB*** | ***ΔfliF*** | ***ΔfliF<br>ΔpleD*** | ***ΔfliF<br>ΔmotB*** | ***ΔflgH*** | ***ΔflgH<br>ΔpleD*** | ***ΔflgH<br>ΔmotB*** | ***ΔflgH<br>ΔpleD<br>ΔmotB*** |
|---|---|---|---|---|---|---|---|---|---|---|---|
| **WT** | X | ns | ** | **** | ns | **** | ns | * | ns | ns | **** |
| ***ΔpleD*** | >0.9999 | X | ** | **** | ns | **** | ns | * | ns | ns | **** |
| ***ΔmotB*** | 0.004 | 0.0039 | X | **** | **** | **** | **** | ns | **** | **** | **** |
| ***ΔpleD<br>ΔmotB*** | <0.0001 | <0.0001 | <0.0001 | X | **** | ns | **** | **** | **** | **** | ** |
| ***ΔfliF*** | 0.5866 | 0.5946 | <0.0001 | <0.0001 | X | **** | ns | **** | ns | ns | **** |
| ***ΔfliF<br>ΔpleD*** | <0.0001 | <0.0001 | <0.0001 | 0.0862 | <0.0001 | X | **** | **** | **** | **** | ns |
| ***ΔfliF<br>ΔmotB*** | 0.584 | 0.5919 | <0.0001 | <0.0001 | >0.9999 | <0.0001 | X | **** | ns | ns | **** |
| ***ΔflgH*** | 0.0127 | 0.0123 | >0.9999 | <0.0001 | <0.0001 | <0.0001 | <0.0001 | X | **** | *** | **** |
| ***ΔflgH<br>ΔpleD*** | 0.0602 | 0.062 | <0.0001 | <0.0001 | 0.9808 | <0.0001 | 0.9812 | <0.0001 | X | ns | **** |
| ***ΔflgH<br>ΔmotB*** | 0.9648 | 0.9668 | <0.0001 | <0.0001 | 0.9993 | <0.0001 | 0.9992 | 0.0002 | 0.6598 | X | **** |
| ***ΔflgH<br>ΔpleD<br>ΔmotB*** | <0.0001 | <0.0001 | <0.0001 | 0.0089 | <0.0001 | 0.999 | <0.0001 | <0.0001 | <0.0001 | <0.0001 | X |

***Table S4*** *Plasmids used in this study*

| **Plasmid** | **Description** | **Antibiotic** | **Reference** |
|---|---|---|---|
| pNPTS138 | Suicide plasmid for making unmarked deletions in *C. crescentus*; carries *sacB* for counter-selection | Km | M. R. Alley |
| pFC1268 | pNPTS138-Δ*flgH*; contains fusion of *CC_2066* flanking regions with first and last 12 nucleotides of *CC_2066* ORF included | Km | (33) |
| pFC3067 | pNPTS138-Δ*pleD*; contains fusion of *CC_2462* flanking regions with first and last 12 nucleotides of *CC_2462* ORF included | Km | (33) |
| pDH543 | pNPTS138-Δ*flmA*; contains fusion of *CC_0233* flanking regions with first and last 12 nucleotides of *CC_0233* ORF included | Km | This work |
| pDH765 | pNPTS138-Δ*motB*; contains fusion of *CC_1573* flanking regions with first and last 12 nucleotides of *CC_1573* ORF included | Km | This work |
| pDH766 | pNPTS138-Δ*fssA*; contains fusion of *CC_1064* flanking regions with first and last 12 nucleotides of *CC_1064* ORF included | Km | This work |
| pDH769 | pNPTS138-Δ*dgcB*; contains fusion of *CC_1850* flanking regions with first and last 12 nucleotides of *CC_1850* ORF included | Km | This work |
| pDH771 | pNPTS138-Δ*shkA*; contains fusion of *CC_0138* flanking regions with first and last 12 nucleotides of *CC_0138* ORF included | Km | This work |
| pDH805 | pNPTS138-Δ*fliF*; contains fusion of *CC_0905* flanking regions with first and last 12 nucleotides of *CC_0905* ORF included | Km | This work |
| pDH810 | pNPTS138-Δ*fliM*; contains fusion of *CC_2061* flanking regions with first and last 12 nucleotides of *CC_2061* ORF included | Km | This work |
| pDH836 | pNPTS138-Δ*fssB*; contains fusion of *CC_2058* flanking regions with first 33 and last 60 nucleotides of *CC_2058* ORF included | Km | This work |
| pMT585 | Contains multiple cloning site downstream of $P_{xyl}$; integrates upstream of *xylX*; used for complementation | Km | (78) |
| pFC3093 | pMT585 containing *flgH* under the control of $P_{flgF}$ for integration at *xylX* locus; 226bp upstream of *CC_2063* fused to the *CC_2066* ORF was inserted in reverse orientation into pMT585 | Km | (33) |
| pFC3094 | pMT585 containing $P_{flgE}$ without insert for integration at *xylX* locus; 216bp upstream of *CC_2063* was inserted in reverse orientation into pMT585 | Km | (33) |
| pDH859 | pMT585 containing $P_{sciP}$ without insert for integration at *xylX* locus; 168bp upstream of *CC_0903* was inserted in reverse orientation into pMT585 | Km | This work |
| pDH879 | pMT585 containing *fliF* under the control of $P_{sciP}$ for integration at *xylX* locus; 168bp upstream of *CC_0903* fused to the *CC_0905* ORF was inserted in reverse orientation into pMT585 | Km | This work |
| pDH852 | pMT585 containing $P_{fliL}$ without insert for integration at *xylX* locus; 226bp upstream of *CC_2062* was inserted in reverse orientation into pMT585 | Km | This work |
| pDH853 | pMT585 containing *fliM* under the control of $P_{fliL}$ for integration at *xylX* locus; 226bp upstream of *CC_2062* fused to the *CC_2061* ORF was inserted in reverse orientation into pMT585 | Km | This work |
| pDH634 | pMT585 containing $P_{flmA}$ without insert for integration at *xylX* locus; 128bp upstream of *CC_0233* was inserted in reverse orientation into pMT585 | Km | This work |
| pDH633 | pMT585 containing *flmA* under the control of $P_{flmA}$ for integration at *xylX* locus; 128bp upstream of *CC_0233* fused to the *CC_0233* ORF was inserted in reverse orientation into pMT585 | Km | This work |

| | | | |
|---|---|---|---|
| pDH893 | pMT585 containing $P_{divK}$ without insert for integration at *xylX* locus; 185bp upstream of *CC_2463* was inserted in reverse orientation into pMT585 | Km | This work |
| pDH880 | pMT585 containing *pleD* under the control of $P_{divK}$ for integration at *xylX* locus; 185bp upstream of *CC_2463* fused to the *CC_2062* ORF was inserted in reverse orientation into pMT585 | Km | This work |
| pDH889 | pMT585 containing $P_{motB}$ without insert for integration at *xylX* locus; 201bp upstream of *CC_1573* was inserted in reverse orientation into pMT585 | Km | This work |
| pDH890 | pMT585 containing *motB* under the control of $P_{motB}$ for integration at *xylX* locus; 201bp upstream of *CC_1573* fused to the *CC_1573* ORF was inserted in reverse orientation into pMT585 | Km | This work |
| pDH935 | pMT585 containing $P_{fssA}$ without insert for integration at *xylX* locus; 68bp upstream of *CC_1064* was inserted in reverse orientation into pMT585 | Km | This work |
| pDH936 | pMT585 containing *fssA* under the control of $P_{fssA}$ for integration at *xylX* locus; 68bp upstream of *CC_1064* fused to the *CC_1064* ORF was inserted in reverse orientation into pMT585 | Km | This work |
| pDH949 | pMT585 containing *fssB* under the control of $P_{fliL}$ for integration at *xylX* locus; 226bp upstream of *CC_2062* fused to the *CC_2058* ORF was inserted in reverse orientation into pMT585 | Km | This work |
| pDH948 | pMT585 containing $P_{dgcB}$ without insert for integration at *xylX* locus; 193bp upstream of *CC_1850* was inserted in reverse orientation into pMT585 | Km | This work |
| pDH937 | pMT585 containing *dgcB* under the control of $P_{dgcB}$ for integration at *xylX* locus; 193bp upstream of *CC_1850* fused to the *CC_1850* ORF was inserted in reverse orientation into pMT585 | Km | This work |
| pDH908 | pMT585 containing $P_{shkA}$ without insert for integration at *xylX* locus; 221bp upstream of *CC_0138* was inserted in reverse orientation into pMT585 | Km | This work |
| pDH909 | pMT585 containing *shkA* under the control of $P_{shkA}$ for integration at *xylX* locus; 221bp upstream of *CC_0138* fused to the *CC_0138* ORF was inserted in reverse orientation into pMT585 | Km | This work |
| pDH904 | To replace *fljK* with $fljK^{T103C}$; pNTPS138 containing $fljK^{T103C}$ allele fused to 488bp upstream and 481 bp downstream flanking regions | Km | This work |
| pFC1948 | pRKlac290 containing the *hfiA* promoter fused to *lacZ* | Tet | (35) |
| pDH967 | pMT585 containing $pleD^{E370Q}$ under the control of $P_{divK}$ for integration at *xylX* locus; 185bp upstream of *CC_2463* fused to the *CC_2062* ORF was inserted in reverse orientation into pMT585 | Km | This work |
| pDH968 | pMT585 containing $motB^{D33N}$ under the control of $P_{motB}$ for integration at *xylX* locus; 201bp upstream of *CC_1573* fused to the *CC_1573* ORF was inserted in reverse oreintation into pMT585 | Km | This work |
| pDH982 | pMT585 containing $pleD^{D53N}$ under the control of $P_{divK}$ for integration at *xylX* locus; 185bp upstream of *CC_2463* fused to the *CC_2062* ORF was inserted in reverse orientation into pMT585 | Km | This work |
| pDH983 | pMT585 containing $dgcB^{E261Q}$ under the control of $P_{dgcB}$ for integration at *xylX* locus; 193bp upstream of *CC_1850* fused to the *CC_1850* ORF was inserted in reverse orientation into pMT585 | Km | This work |

***Table S5*** *Strains used in this study*

| Strain | Organism | Genotype | Description | Source |
|---|---|---|---|---|
| FC19 | *C. crescentus* CB15 | CB15 | Wild type | ATCC 19089 |
| FC1266 | *C. crescentus* CB15 | Δ*flgH* | In frame deletion of *CC_2066* | (33) |
| DH816 | *C. crescentus* CB15 | Δ*fliF* | In frame deletion of *CC_0905* | This work |
| DH843 | *C. crescentus* CB15 | Δ*fliF* Δ*pleD* | In frame deletion of *CC_0905* in DH659 background | This work |
| DH844 | *C. crescentus* CB15 | Δ*fliF* Δ*motB* | In frame deletion of *CC_0905* in DH776 background | This work |
| DH839 | *C. crescentus* CB15 | Δ*fliM* | In frame deletion of *CC_2061* | This work |
| DH858 | *C. crescentus* CB15 | Δ*fliM* Δ*motB* | In frame deletion of *CC_2061* in DH776 background | This work |
| DH840 | *C. crescentus* CB15 | Δ*fliM* Δ*pleD* | In frame deletion of *CC_2061* in DH659 background | This work |
| DH777 | *C. crescentus* CB15 | Δ*flgH* Δ*motB* | In frame deletion of *CC_1573* in FC1266 background | This work |
| DH649 | *C. crescentus* CB15 | Δ*flgH* Δ*pleD* | In frame deletion of *CC_2462* in FC1266 background | (33) |
| DH547 | *C. crescentus* CB15 | Δ*flmA* | In frame deletion of *CC_0233* | This work |
| DH851 | *C. crescentus* CB15 | Δ*flmA* Δ*motB* | In frame deletion of *CC_0233* in DH776 background | This work |
| DH850 | *C. crescentus* CB15 | Δ*flmA* Δ*pleD* | In frame deletion of *CC_0233* in DH659 background | This work |
| DH659 | *C. crescentus* CB15 | Δ*pleD* | In frame deletion of *CC_2462* | (33) |
| DH776 | *C. crescentus* CB15 | Δ*motB* | In frame deletion of *CC_1573* | This work |
| DH783 | *C. crescentus* CB15 | Δ*fssA* | In frame deletion of *CC_1064* | This work |
| DH784 | *C. crescentus* CB15 | Δ*fssA* Δ*flgH* | In frame deletion of *CC_1064* in FC1266 background | This work |
| DH875 | *C. crescentus* CB15 | Δ*fssA* Δ*fliF* | In frame deletion of *CC_1064* in DH816 background | This work |
| DH845 | *C. crescentus* CB15 | Δ*fssB* | In frame deletion of *CC_2058* | This work |
| DH846 | *C. crescentus* CB15 | Δ*fssB* Δ*flgH* | In frame deletion of *CC_2058* in FC1266 background | This work |
| DH878 | *C. crescentus* CB15 | Δ*fssB* Δ*fliF* | In frame deletion of *CC_2058* in DH816 background | This work |
| DH787 | *C. crescentus* CB15 | Δ*dgcB* | In frame deletion of *CC_1850* | This work |
| DH788 | *C. crescentus* CB15 | Δ*dgcB* Δ*flgH* | In frame deletion of *CC_1850* in FC1266 background | This work |
| DH876 | *C. crescentus* CB15 | Δ*dgcB* Δ*fliF* | In frame deletion of *CC_1850* in DH816 background | This work |
| DH791 | *C. crescentus* CB15 | Δ*shkA* | In frame deletion of *CC_0138* | This work |
| DH792 | *C. crescentus* CB15 | Δ*shkA* Δ*flgH* | In frame deletion of *CC_0138* in FC1266 background | This work |
| DH897 | *C. crescentus* CB15 | Δ*shkA* Δ*fliF* | In frame deletion of *CC_0138* in DH816 background | This work |
| DH911 | *C. crescentus* CB15 | $fljK^{T103C}$ | $fljK^{T103C}$ inserted at *fljK* locus using pDH904 | This work |
| DH912 | *C. crescentus* CB15 | Δ*flgH* $fljK^{T103C}$ | $fljK^{T103C}$ inserted at *fljK* locus of DH553 using pDH904 | This work |

| | | | | |
|---|---|---|---|---|
| DH946 | *C. crescentus* CB15 | Δ*fssA* $fljK^{T103C}$ | $fljK^{T103C}$ inserted at *fljK* locus of DH783 using pDH904 | This work |
| DH947 | *C. crescentus* CB15 | Δ*fssB* $fljK^{T103C}$ | $fljK^{T103C}$ inserted at *fljK* locus of DH845 using pDH904 | This work |
| DH913 | *C. crescentus* CB15 | Δ*pleD* Δ*motB* | In frame deletion of *CC_1573* in DH659 background | This work |
| DH914 | *C. crescentus* CB15 | Δ*flgH* Δ*pleD* Δ*motB* | In frame deletion of *CC_1573* in DH649 background | This work |
| DH822 | *C. crescentus* CB15 | Δ*fliF* Δ*flgH* | In frame deletion of *CC_0905* in FC1266 background | This work |
| DH964 | *C. crescentus* CB15 | Δ*fliF* Δ*flgH* Δ*motB* | In frame deletion of *CC_2066* in DH844 background | This work |
| DH996 | *C. crescentus* CB15 | Δ*fliF* Δ*flgH* Δ*pleD* | In frame deletion of *CC_2462* in DH822 background | This work |
| FC3075 | *C. crescentus* CB15 | Δ*flgH* *xyl*::$P_{flgE}$-*empty* | pFC3094 integrated at xylose locus of FC1266 | (33) |
| FC3074 | *C. crescentus* CB15 | Δ*flgH* *xyl*::$P_{flgE}$-*flgH* | pFC3093 integrated at xylose locus of FC1266 | (33) |
| DH887 | *C. crescentus* CB15 | Δ*fliF* *xyl*::$P_{sciP}$-*empty* | pDH859 integrated at xylose locus of DH816 | This work |
| DH888 | *C. crescentus* CB15 | Δ*fliF* *xyl*::$P_{sciP}$-*fliF* | pDH879 integrated at xylose locus of DH816 | This work |
| DH861 | *C. crescentus* CB15 | Δ*fliM* *xyl*::$P_{fliL}$-*empty* | pDH852 integrated at xylose locus of DH839 | This work |
| DH862 | *C. crescentus* CB15 | Δ*fliM* *xyl*::$P_{fliL}$-*fliM* | pDH853 integrated at xylose locus of DH839 | This work |
| DH638 | *C. crescentus* CB15 | Δ*flmA* *xyl*::$P_{flmA}$-*empty* | pDH634 integrated at xylose locus of DH547 | This work |
| DH637 | *C. crescentus* CB15 | Δ*flmA* *xyl*::$P_{flmA}$-*flmA* | pDH633 integrated at xylose locus of DH547 | This work |
| DH899 | *C. crescentus* CB15 | Δ*pleD* *xyl*::$P_{divK}$-*empty* | pDH893 integrated at xylose locus of DH659 | This work |
| DH898 | *C. crescentus* CB15 | Δ*pleD* *xyl*::$P_{divK}$-*pleD* | pDH880 integrated at xylose locus of DH659 | This work |
| DH906 | *C. crescentus* CB15 | Δ*pleD* Δ*flgH* *xyl*::$P_{divK}$-*empty* | pDH893 integrated at xylose locus of DH649 | This work |
| DH905 | *C. crescentus* CB15 | Δ*pleD* Δ*flgH* *xyl*::$P_{divK}$-*pleD* | pDH880 integrated at xylose locus of DH649 | This work |
| DH895 | *C. crescentus* CB15 | Δ*motB* *xyl*::$P_{motB}$-*empty* | pDH889 integrated at xylose locus of DH776 | This work |
| DH896 | *C. crescentus* CB15 | Δ*motB* *xyl*::$P_{motB}$-*motB* | pDH890 integrated at xylose locus of DH776 | This work |
| DH902 | *C. crescentus* CB15 | Δ*motB* Δ*flgH* *xyl*::$P_{motB}$-*empty* | pDH889 integrated at xylose locus of DH777 | This work |
| DH903 | *C. crescentus* CB15 | Δ*motB* Δ*flgH* *xyl*::$P_{motB}$-*motB* | pDH890 integrated at xylose locus of DH777 | This work |
| DH938 | *C. crescentus* CB15 | Δ*fssA* *xyl*::$P_{fssA}$-*empty* | pDH935 integrated at xylose locus of DH783 | This work |
| DH939 | *C. crescentus* CB15 | Δ*fssA* *xyl*::$P_{fssA}$-*fssA* | pDH936 integrated at xylose locus of DH783 | This work |

| | | | | |
|---|---|---|---|---|
| DH940 | *C. crescentus* CB15 | *ΔfssA ΔflgH xyl::$P_{fssA}$-empty* | pDH935 integrated at xylose locus of DH784 | This work |
| DH941 | *C. crescentus* CB15 | *ΔfssA ΔflgH xyl::$P_{fssA}$-fssA* | pDH936 integrated at xylose locus of DH784 | This work |
| DH950 | *C. crescentus* CB15 | *ΔdgcB xyl::$P_{dgcB}$-empty* | pDH948 integrated at xylose locus of DH787 | This work |
| DH951 | *C. crescentus* CB15 | *ΔdgcB xyl::$P_{dgcB}$-dgcB* | pDH937 integrated at xylose locus of DH787 | This work |
| DH952 | *C. crescentus* CB15 | *ΔdgcB ΔflgH xyl::$P_{dgcB}$-empty* | pDH948 integrated at xylose locus of DH788 | This work |
| DH953 | *C. crescentus* CB15 | *ΔdgcB ΔflgH xyl::$P_{dgcB}$-dgcB* | pDH937 integrated at xylose locus of DH788 | This work |
| DH954 | *C. crescentus* CB15 | *ΔdgcB ΔshkA* | In frame deletion of *CC_0138* in DH787 background | This work |
| DH955 | *C. crescentus* CB15 | *ΔflgH ΔdgcB ΔshkA* | In frame deletion of *CC_0138* in DH788 background | This work |
| DH957 | *C. crescentus* CB15 | *ΔfssB xyl::$P_{fliL}$-empty* | pDH852 integrated at xylose locus of DH845 | This work |
| DH958 | *C. crescentus* CB15 | *ΔfssB xyl::$P_{fliL}$-fssB* | pDH949 integrated at xylose locus of DH845 | This work |
| DH959 | *C. crescentus* CB15 | *ΔfssB ΔflgH xyl::$P_{fliL}$-empty* | pDH852 integrated at xylose locus of DH846 | This work |
| DH960 | *C. crescentus* CB15 | *ΔfssB ΔflgH xyl::$P_{fliL}$-fssB* | pDH949 integrated at xylose locus of DH846 | This work |
| DH920 | *C. crescentus* CB15 | *ΔshkA xyl::$P_{shkA}$-empty* | pDH908 integrated at xylose locus of DH791 | This work |
| DH921 | *C. crescentus* CB15 | *ΔshkA xyl::$P_{shkA}$-shkA* | pDH909 integrated at xylose locus of DH791 | This work |
| DH922 | *C. crescentus* CB15 | *ΔshkA ΔflgH xyl::$P_{shkA}$-empty* | pDH908 integrated at xylose locus of DH792 | This work |
| DH923 | *C. crescentus* CB15 | *ΔshkA ΔflgH xyl::$P_{shkA}$-shkA* | pDH909 integrated at xylose locus of DH792 | This work |
| DH990 | *C. crescentus* CB15 | *ΔmotB ΔflgH xyl::$P_{motB}$-motB$^{D33N}$* | pDH968 integrated at x12ylose locus of DH777 | This work |
| DH991 | *C. crescentus* CB15 | *ΔdgcB ΔflgH xyl::$P_{dgcB}$-dgcB$^{E271Q}$* | pDH983 integrated at x12ylose locus of DH788 | This work |
| DH997 | *C. crescentus* CB15 | *ΔpleD ΔflgH xyl::$P_{divK}$-pleD$^{E370Q}$* | pDH967 integrated at 12xylose locus of DH649 | This work |
| DH998 | *C. crescentus* CB15 | *ΔpleD ΔflgH xyl::$P_{divK}$-pleD$^{D53N}$* | pDH982 integrated at 12xylose locus of DH649 | This work |
| DH1037 | *C. crescentus* CB15 | *ΔmotA* | In frame deletion of *CC_0750* | This work |
| DH1040 | *C. crescentus* CB15 | *ΔfliF ΔmotA* | In frame deletion of *CC_0750* in DH816 background | This work |
| DH1041 | *C. crescentus* CB15 | *ΔflgH ΔmotA* | In frame deletion of *CC_0750* in DH553 | This work |
| DH1086 | *C. crescentus* CB15 | *ΔfssA ΔfssB* | In frame deletion of *CC_1064* in DH845 background | This work |
| DH1086 | *C. crescentus* CB15 | *ΔflgH ΔfssA ΔfssB* | In frame deletion of *CC_1064* in DH846 background | This work |
| DH1098 | *C. crescentus* CB15 | *ΔshkA ΔmotB* | In frame deletion of *CC_1573* in DH791 background | This work |

| | | | | |
|---|---|---|---|---|
| DH1099 | *C. crescentus* CB15 | Δ*flgH* Δ*shkA* Δ*motB* | In frame deletion of *CC_1573* in DH792 background | This work |

***Table S6*** *Features of the barcoded transposon libraries used in this study*

| **Library name** | **SCD_147** | **SCD_105** |
|---|---|---|
| SRA BioProject number | PRJNA640825 | PRJNA640725 |
| Genetic background | CB15 WT | Δ*flgH* |
| Total reads | 62819957 | 48083939 |
| Mapped reads | 35866380 | 23139968 |
| Mapped barcodes in pool | 273529 | 323456 |
| Distinct insertion sites | 62467 | 65051 |
| Coding genes with central insertions | 3244 | 3224 |
| Median strains per hit protein | 40 | 47 |
| Mean strains per hit protein | 58.2 | 69.4 |

***Figure S1*** *Surface contact independent activation of holdfast production in* ΔflgH A) Shedding of holdfast polysaccharide into spent medium. Δ*flgH* releases a holdfast specific fWGA reactive material into the spent medium during growth in M2X liquid. B) Comparison showing the fraction of holdfast producing cells in wild type and Δ*flgH* backgrounds. Centrifugation has no effect on holdfast production in either strain. C) Micrographs of wild type and Δ*flgH* mutant cells taken immediately after direct staining of holdfast in liquid cultures.

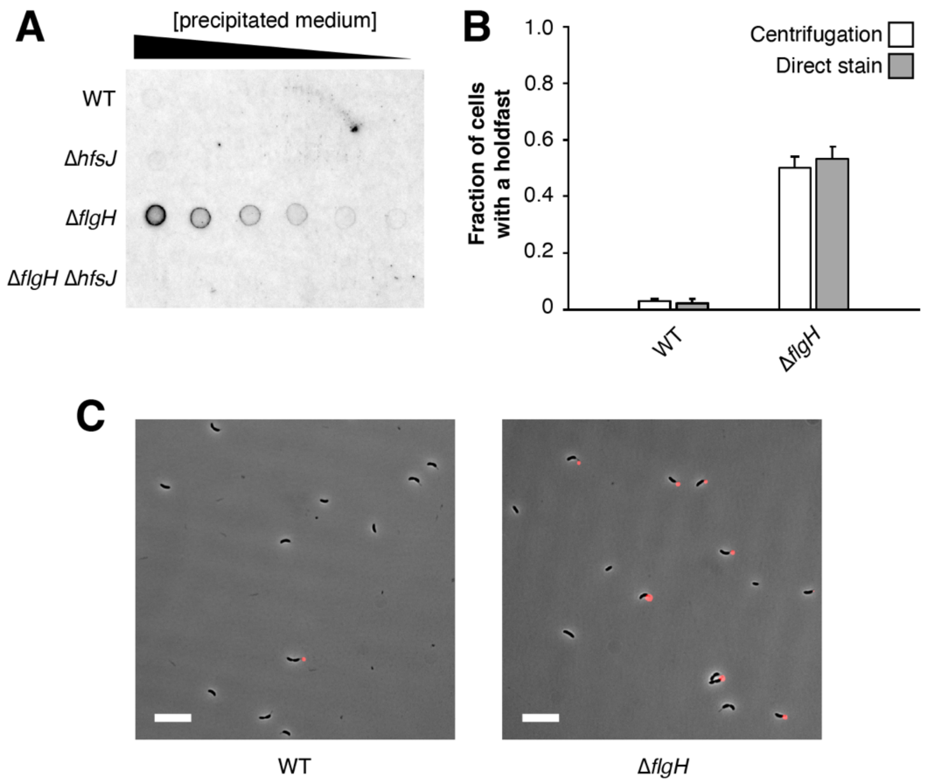

***Figure S2*** *Confirmation of the two-pathway model for flagellar control of holdfast production* A) CV staining of mutants from Fig2D grown in PYE. The loss of adhesion when the mechanical (Δ*motB*) and developmental (Δ*pleD*) pathways are inactivated simultaneously is not specific to defined M2X medium. Mean values from six biological replicates are shown along with their associated standard deviations. B) CV stain confirming the placement of *motA* in the mechanical pathway. Δ*motA* reduces hyper-adhesion specifically in late flagellar mutants, mirroring the Δ*motB* suppression pattern. Mean values from seven biological replicates are shown along with their associated standard deviations. C) CV stain showing additive effects of disrupting *shkA* and *motB* simultaneously. The results confirm the placement of *shkA* and *motB* in separate signaling pathways. Mean values from six biological replicates are shown along with their associated standard deviations. D) CV stain showing additive effects of disrupting *shkA* and *dgcB* simultaneously. The results confirm the placement of *shkA* and *dgcB* in separate signaling pathways. Mean values from six biological replicates are shown along with their associated standard deviations.

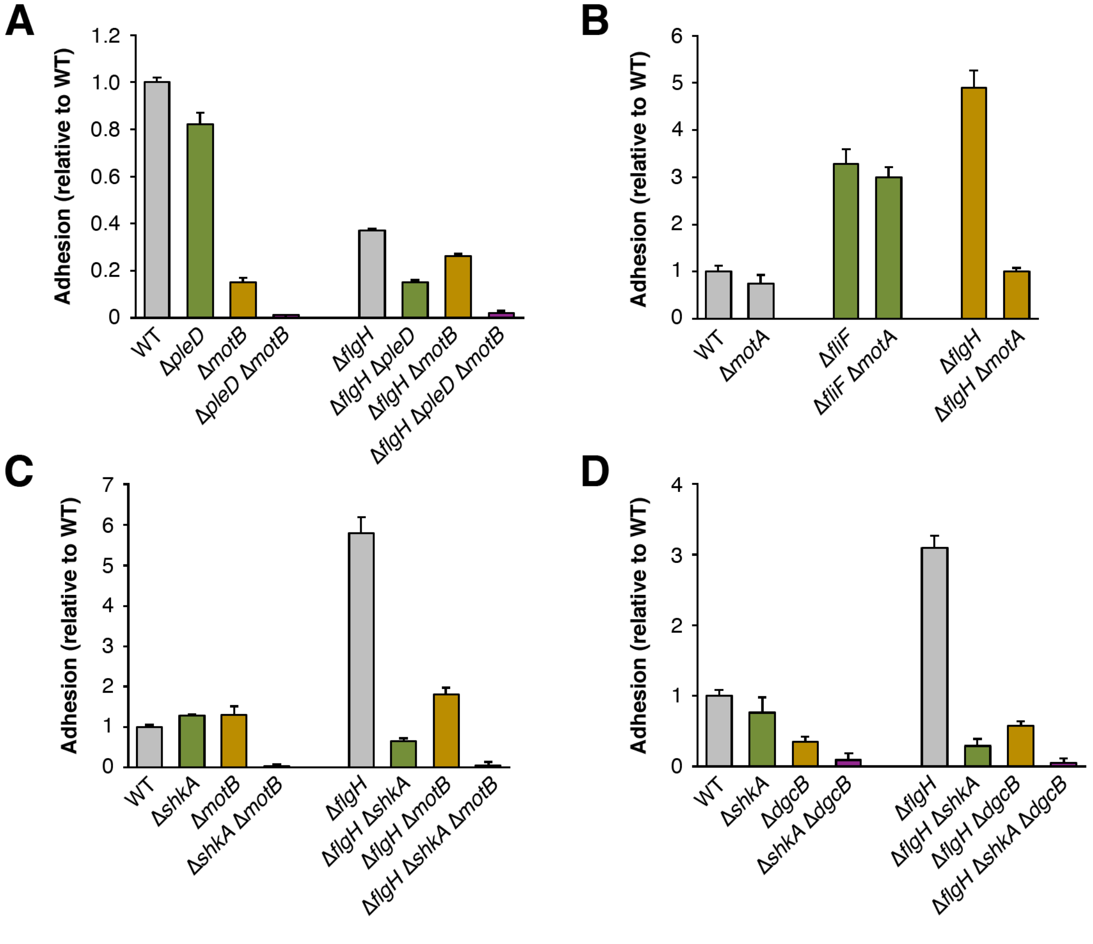

***Figure S3*** *Additional analysis of the* Δ*fssA phenotype* A) Mapping of mutations that suppress the Δ*fssA* motility phenotype by whole genome sequencing. The suppressor numbers correspond to files in SRA accession PRJNA672134. Nucleotide positions correspond to coordinates in the NA1000 genome. B) Soft-agar motility assay showing that the Δ*fssB* motility defect is epistatic to the Δ*fssA* phenotype. Motile suppressors do not appear in the Δ*fssA* Δ*fssB* double mutant after a 96 hour incubation. C) CV staining experiment showing that the Δ*fssA* and Δ*fssB* are not additive for suppression of hyper-adhesion in the Δ*flgH* mutant. Mean values from six biological replicates are shown along with their associated standard deviations.

**A**

| Suppressor | Position | Mutation | Substitution |
|---|---|---|---|
| 1 | 1,739,423 | T=>G | MotB S52R |
| 2 | 825,347 | A=>G | MotA F15L |
| 3 | 1,739,423 | T=>A | MotB S52C |
| 4 | 824,839 | A=>T | MotA L184Q |
| 5 | 1,739,422 | C=>T | MotB S52N |
| 6 | 1,739,423 | T=>A | MotB S52C |
| 7 | 824,839 | A=>T | MotA L184Q |
| 8 | 1,739,434 | A=>C | MotB L48R |
| 9 | 1,739,393 | A=>G | MotB Y62H |
| 10 | 1,739,423 | T=>A | MotB S52C |
| 11 | 1,739,423 | T=>A | MotB S52C |
| 12 | 1,739,423 | T=>A | MotB S52C |
| 13 | 1,739,423 | T=>C | MotB S52G |
| 14 | 1,739,423 | T=>A | MotB S52C |

**B**

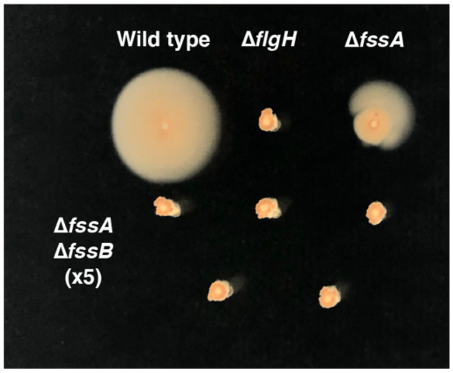

**C**

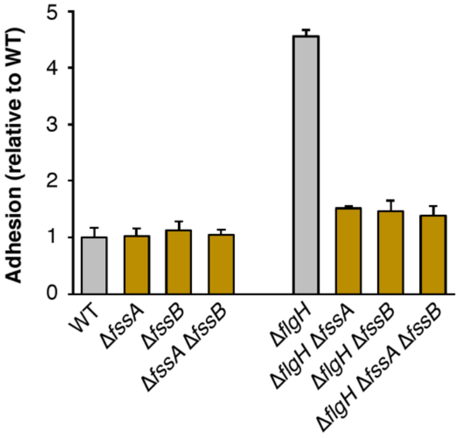

***Figure S4*** *Genetic complementation of surface attachment and motility phenotypes* A) Complementation of flagellar hierarchy mutants in soft agar. B) Complementation of hyper-adhesion in flagellar mutants. C) Complementation of crystal violet staining for *fss* mutants in Δ*flgH* background. D, E) Restoration of motility phenotypes in soft agar for *fss* mutants.

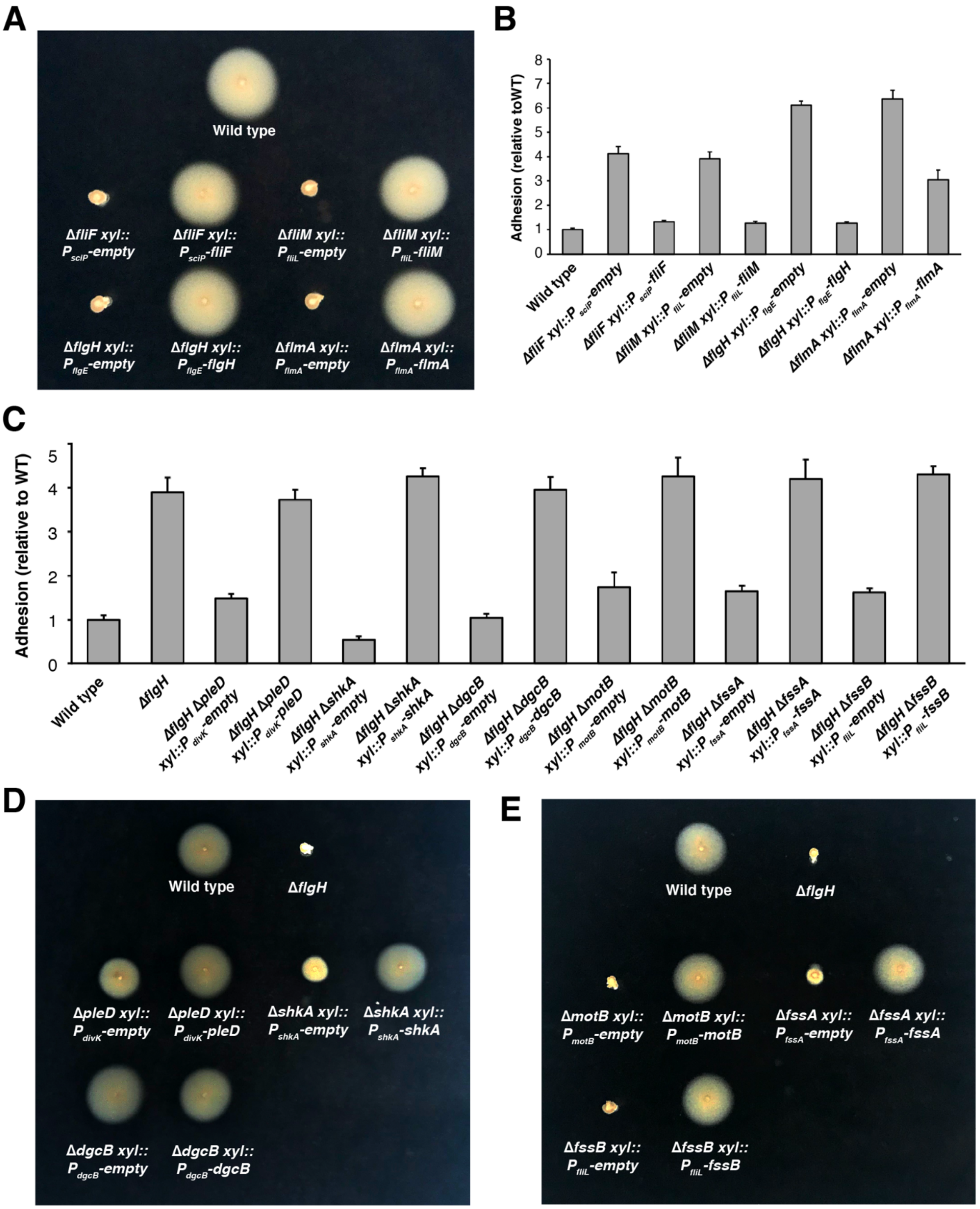
